# High-resolution analysis of individual *Drosophila melanogaster* larvae within groups uncovers inter- and intra-individual variability in locomotion and its neurogenetic modulation

**DOI:** 10.1101/2022.09.27.509663

**Authors:** Michael Thane, Emmanouil Paisios, Torsten Stöter, Anna-Rosa Krüger, Sebastian Gläß, Anne-Kristin Dahse, Nicole Scholz, Bertram Gerber, Dirk J. Lehmann, Michael Schleyer

## Abstract

Neuronally orchestrated muscular movement and locomotion are defining faculties of multicellular animals. Due to its numerically simple brain and neuromuscular system and its genetic accessibility, the larva of the fruit fly *Drosophila melanogaster* is an established model to study these processes at tractable levels of complexity. However, although the faculty of locomotion clearly pertains to the individual animal, present studies of locomotion in larval *Drosophila* mostly use group assays and measurements aggregated across individual animals. The alternative is to measure animals one at a time, an extravagance for larger-scale analyses. In principle or in practice, this in particular rules out grasping the inter- and intra-individual variability in locomotion and its genetic and neuronal determinants. Here we present the IMBA (Individual Maggot Behaviour Analyser) for tracking and analysing the behaviour of individual larvae within groups. Using a combination of computational modelling and statistical approaches, the IMBA reliably resolves individual identity across collisions. It does not require specific hardware and can therefore be used in non-expert labs. We take advantage of the IMBA first to systematically describe the inter- and intra-individual variability in free, unconstrained locomotion in wild-type animals. We then report the discovery of a novel, complex locomotion phenotype of a mutant lacking an adhesion-type GPCR. The IMBA further allows us to determine, at the level of individual animals, the modulation of locomotion across repeated activations of dopamine neurons. Strikingly, IMBA can also be used to analyse ‘silly walks’, that is patterns of locomotion it was not originally designed to investigate. This is shown for the transient backward locomotion induced by brief optogenetic activation of the brain-descending ‘mooncrawler’ neurons, and the variability in this behaviour. Thus, the IMBA is an easy-to-use toolbox allowing an unprecedentedly rich view of the behaviour and behavioural variability of individual *Drosophila* larvae, with utility in multiple biomedical research contexts.

## 1 Introduction

Understanding the neuronal control of movement and locomotion, that is of how we do what we do, is an important and rewarding task for basic research in the behavioural and neural sciences. Furthermore, such an understanding can inspire the development of medical and technical applications, for example regarding gait analysis for medical diagnosis, or bio-inspired robotics [1–3]. A comprehensive analysis of neuromuscular control in humans remains a formidable challenge, however, not least because of the complexity of our brains and bodies, complexity that is scarcely less in other mammals. In the past few years, the larva of the fruit fly *Drosophila melanogaster* has emerged as a model organism for studying many aspects of these processes [4]. The combination of a rich toolbox for cell-specific transgenic manipulation, a nervous system of only about 10,000 neurons, and the availability of an electron-microscope-based synaptic connectome allows neuronal processes to be studied in unrivalled detail. The utility of the larva for behavioural neuroscience has been further aided both by its relatively simple body and the relatively moderate level of complexity in its locomotion. Locomotion in *Drosophila* larvae can, simplifying somewhat, be described as consisting of periods of relatively straight runs characterized by peristaltic waves of muscle contraction along the body wall that alternate with occasional lateral head movements [4–14]. This results in a zig-zagging pattern of locomotion towards their target, similarly observed in adult flies and mosquitoes.

Most studies of locomotion in larval *Drosophila* either use *en-masse*, group assays and measurements aggregated across individual animals, or they need to measure single animals one at a time, which practically precludes larger-scale analyses. It has thus remained difficult to come to grips with the causes and consequences of the variability in locomotion both between and within individual animals (henceforth called inter-individual and intra-individual variability, respectively). Yet individual idiosyncrasies in behaviour are described across the animal kingdom, including *Drosophila* [15, 16]. Such inter-individual variance in behaviour may be regarded not as noise, but as an inroad to studying the determinants and implications of such variability and (if these are heritable) as raw material for natural selection. Likewise, intra-individual variability in behaviour is essential to initiate operant learning, that is to generate novel actions capable of bringing about desired outcomes. To study behavioural differences between and across individuals efficiently, we need to be able to (1) observe behaviour in individual animals, (2) measure large numbers of animals in a short period of time, and (3) manipulate the candidate cellular and molecular substrates of the observed behaviour easily and precisely.

An increasing number of video-tracking approaches have been developed in the past decade to facilitate such analyses in larval *Drosophila*. Some of these were designed to track the behaviour of individually assayed animals, e.g. [17–20]. These usually provide a very high temporal and spatial resolution, but are extravagantly time-consuming if high sample sizes are required. Other approaches were designed to track groups of animals, e.g. [21–28]. These focus on the average behaviour of groups of animals and thus do not allow for individuals to be followed. More recent approaches have tried to combine the advantages of both types of tracking [29, 30]. The major challenge in these cases is that individual *Drosophila* larvae look very much alike. As a result, when two larvae collide, their identity is lost, resulting in a multitude of separated tracks and no opportunity for the experimenter to disentangle which tracks belong to which of the animals. Two different strategies have been followed in tackling this problem: one relies on the parallel tracking of many separated individual animals [29]; the other tries to resolve the collisions between larvae and thus to keep track of the identity of individual larvae within a group of animals [30] (for an earlier approach not specific to *Drosophila* larvae, see [31]). The latter solution has been integrated into the FIM (Frustrated total internal reflection-based Imaging Method) tracker software which requires specific hardware to illuminate the tracked animals [23]. Although both these approaches are suitable to track and analyse high numbers of individual animals, they both have specific requirements regarding the physical setup for video-recording, and therefore require extensive redesigns of established behavioural assays in order to be used. We follow a different approach by starting from an established standard behavioural assay and developing a tracking solution that works for that assay and that non-expert users can easily implement in their research programs using that assay or its derivatives.

Here we present the IMBA, the Individual Maggot Behaviour Analyser. It consists of the IMBAtracker and the IMBAvisualiser. Together, these can track and analyse the behaviour of individual larvae within groups. They allow collisions to be resolved between individual larvae using a combination of statistical and computational approaches that can be applied to videos obtained by non-expert users in standard experimental assays. They do not require any specific hardware beyond a camera and stable lighting, and can therefore be easily applied to various derivatives of these standard behavioural assays, including ones that are commonly used for olfactory, gustatory or visual choice [32–36], or associative learning [37–39]. We further show the utility of the IMBA as a research tool by discovering a novel, complex locomotion phenotype in mutants of an adhesion G-protein coupled receptor gene (*Cirl*) [40] and by studying the effects of repeatedly activating dopamine neurons (as covered by the *TH-Gal4* driver) on locomotion. The utility of the IMBA is further demonstrated for patterns of locomotion it was not originally designed to study, using as an example the backward movements induced by optogenetic activation of the so-called ‘mooncrawler’ neurons (as covered by the *R53F07-Gal4* driver). In all these cases we pay particular attention to the inter-individual and intra-individual variability in behaviour.

## 2 Results

In this study, we investigate the behaviour of individuals within groups of *Drosophila* larvae. Our approach to the problem of losing track of individual identities upon collisions involves two critical steps: first, for our analysis we only considered tracks that were more than half the length of the recorded video, so we were sure that no two tracks in our analysis belonged to the same individual animal; second, we tried to stitch together tracks from the same individual across collisions in order to maximize the number of individual animals that we could include within our analysis. To this end, we employed a combination of a computational approach modelling the shape of larvae and keeping their identities during a collision, and a statistical approach comparing larvae before and after a collision. For more details, see the Materials and Methods section.

### 2.1 The IMBA allows the behaviour of individuals within groups to be studied

In order to evaluate our approach, we compared the results with manual assessments of collisions by an experienced experimenter. In 48 videos with approximately 20 larvae each, 1844 collisions were found by the experimenter. We attempted to resolve 1531 by means of the IMBAtracker, and of these, 1511 were correctly resolved (Fig. 1-E1A). In total, 83.7 % of all tracked data belonged to tracks that fulfilled our criterion of being at least half the length of the video after collision resolution and were used in the analysis (Fig. 1-E1B-D).

In the next step, we developed an R-based shiny application to analyse and visualize the behaviour of these individual larvae. This application, called the IMBAvisualiser, calculates a total of 95 attributes for each animal that can be explored and visualized in an interactive manner. For an overview of the process, see Figure 1-E2. Both the code and a full documentation of this application can be accessed via GitHub at https://github.com/XXX and will be regularly updated and further developed in the future.

The locomotion of *Drosophila* larvae has been studied in detail in the past decade [5–7]. In brief, the locomotion can be described as sequences of run phases, consisting of relatively straight, forward peristaltic movement, and reorientation manoeuvres, consisting of lateral head casts (HC), which are usually associated with a strong reduction in speed and are followed by changes in direction (Fig. 1A) [8]. Accordingly, we first defined HC phases by the above-threshold angular velocity of the animal’s head vector (Fig. 1B-C) (for details, see the Materials and Methods section). Animals were considered to be in a run phase whenever they were not in a HC phase. A period of 1.5 s before and after each HC was excluded from the run phases to prevent the potential effects of HCs on run behaviour. All these definitions are in line with our previous work [27, 41]. For a more detailed analysis of run periods, we defined the cycles of peristaltic movement by means of the animal’s forward velocity (Fig. 1D). Each local maximum of forward velocity during a run was defined as one “step”, with the period between two steps (inter-step, IS) capturing one complete cycle of peristaltic movement (for a similar approach, see [42]).

**Fig. 1:**
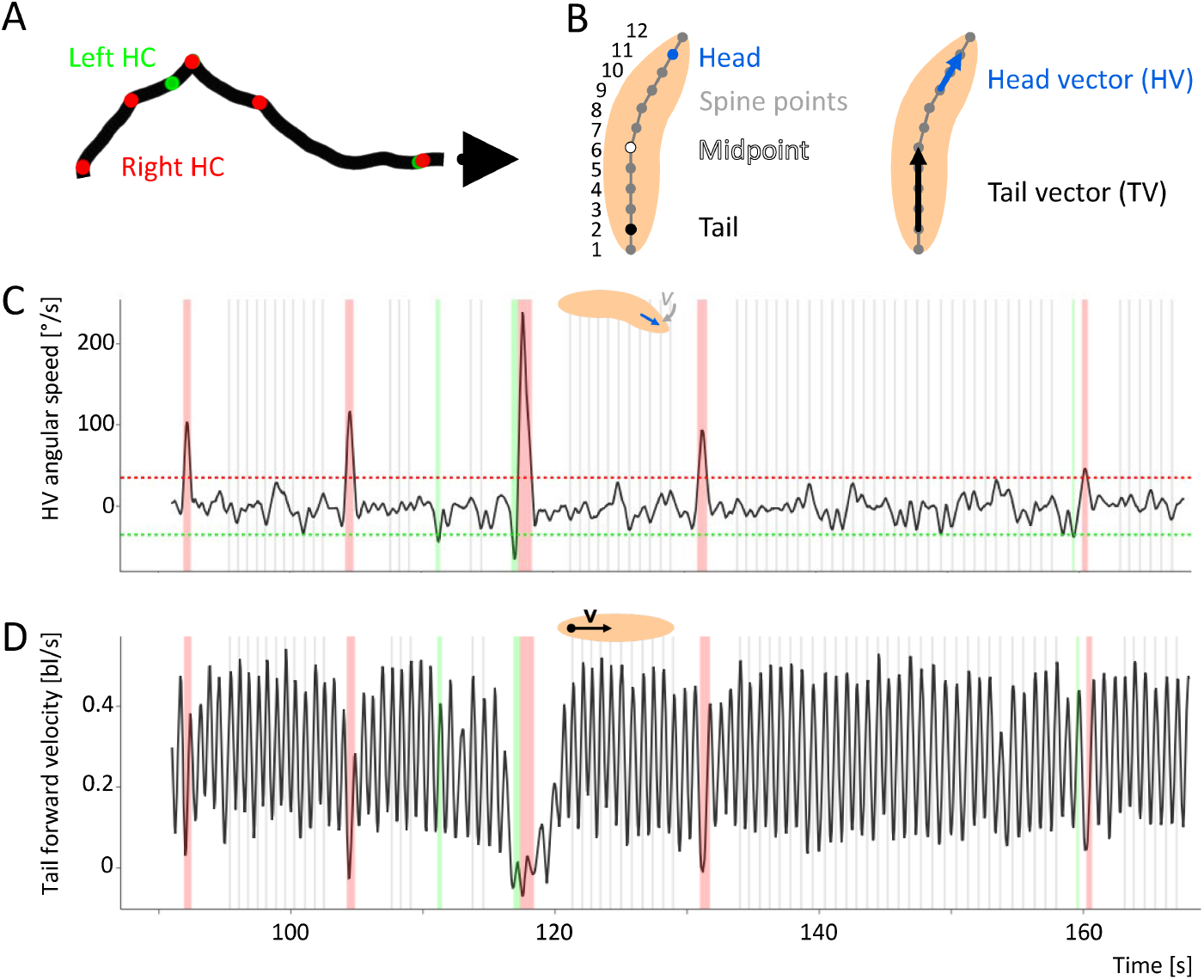
Principles of the analysis. (A) The track of the midpoint of a sample larva. Green and red dots indicate left and right HCs, respectively. The arrow head indicates the end of the track. (B) (left) 12 equidistant spine points are determined along the spine of each larva, including the back and front “tip” (spine points 1 and 12, respectively). The second spine point from the front (i.e. number 11) is taken as indicative of the head, the second point from the rear as indicative of the tail (i.e. number 2). (right) A head vector (HV) is determined from spine points 9 to 11, and a tail vector (TV) from spine points 2 to 6. (C) The angular speed of the head vector of the sample larva displayed in (A). Negative values indicate movement to the left, positive values movement to the right. A HC is detected when the HV angular speed exceeds a threshold of ±35 °/s (stippled lines). In this and the following figures, left and right HCs are indicated by green and red shadings, respectively. (D) The forward velocity of the tail of the sample larva displayed in (A). During runs, the velocity oscillates regularly. Each oscillation marks one cycle of peristaltic forward movement. Therefore, we define each peak of the Tail forward velocity during runs as one “step”, indicated by grey lines in this and analogous figures. During HCs, the regular forward movement is stopped. Therefore, 1.5 s before and after each HC, no steps are detected.

### 2.2 Attributes of basic locomotion in larvae are variable across individuals

On the basis of these observations, and considering the behaviour of 875 innate, i.e. experimentally naïve, animals, we determined a set of six behavioural attributes to describe the basic locomotion of larvae: the IS speed, indicating the average speed across all peristaltic cycles (Fig. 2A); the IS distance, indicating the average size of the steps (Fig. 2B); the IS interval, indicating the average time between two steps (Fig. 2C); the absolute bending angle, indicating how much the body of a larva was bent throughout the whole observation period (Fig. 2D); the HC rate, indicating how many HCs an animal performed per second (Fig. 2E); and the absolute HC angle, indicating the average size of an animal’s HCs (Fig. 2F). To determine the variability in each behavioural attribute, we calculated the coefficient of variation (CV), which indicates the standard deviation as a percentage of the mean and allows the inter-individual variability to be compared across different behavioural attributes. The CV was lowest for the IS distance (15.6 %) and highest for the HC rate (47.4 %).

**Fig. 2:**
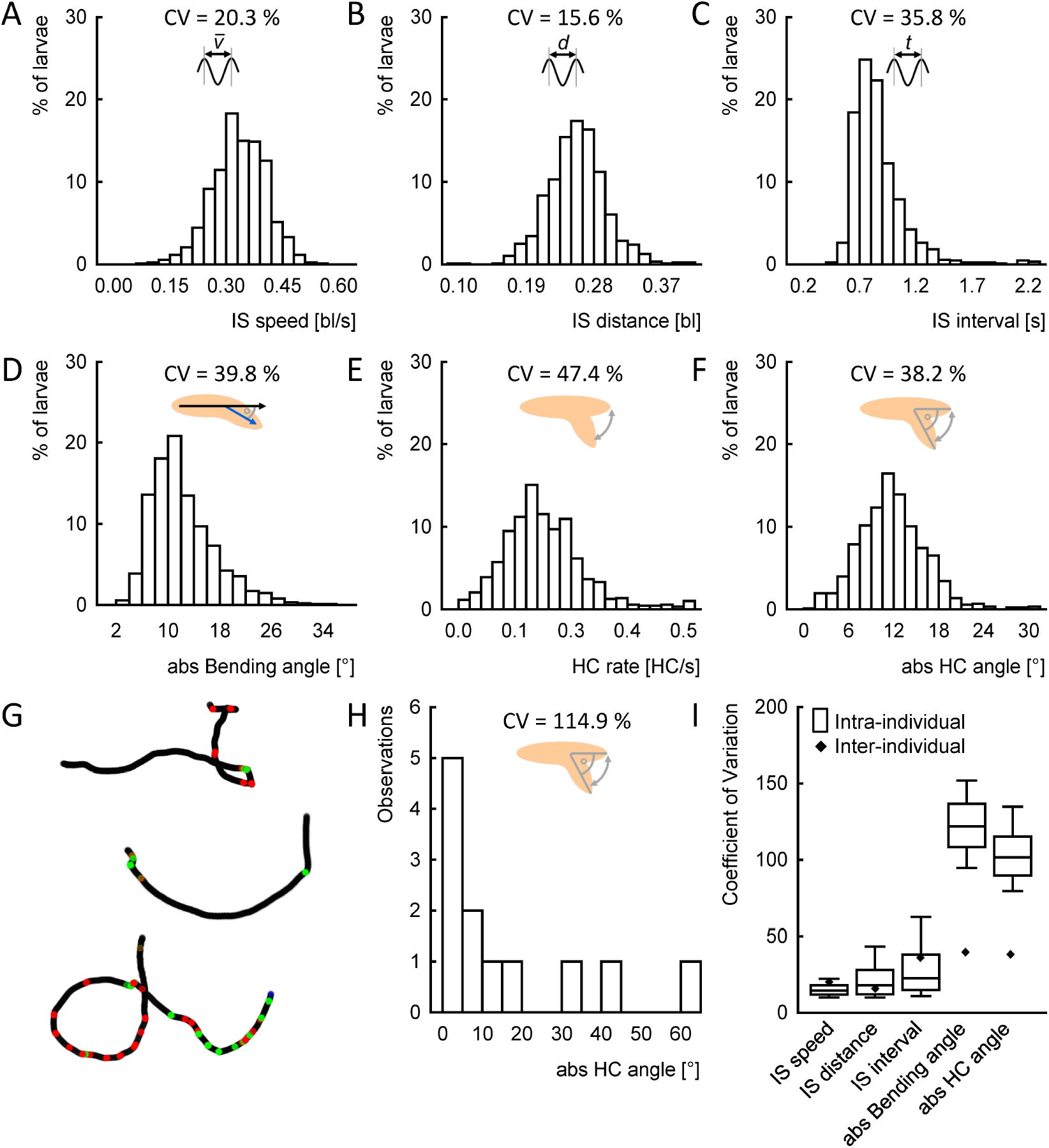
Variability of basic locomotor attributes of individual *Drosophila* larvae. Six basic locomotor attributes were determined: (A) the inter-step (IS) speed, i.e. the average speed of each larva’s midpoint during runs, (B) the IS distance, i.e. the average distance travelled within one peristaltic cycle for each larva, (C) the IS interval, i.e. the average time required for one peristaltic cycle for each larva, (D) the absolute bending angle, i.e. how much each larva was bent throughout the observation period, (E) the HC rate, i.e. the number of HCs per second of each larva, and (F) the absolute HC angle, measuring the average size of each larva’s HCs. Displayed are histograms of each behavioural attribute per individual, together with the coefficient of variation (CV), which indicates the standard deviation of the attribute, divided by its mean, across all individuals. (G) Three sample tracks of individuals showing different patterns of behaviour: (top) a stretch of relatively straight, forward locomotion, followed by a series of rather large HCs and turns, (middle) a stretch of curved forward locomotion, flanked by HCs, (bottom) continuous, small HCs. (H) Absolute HC angles of the top sample individual from (G) range from close to zero up to 60°. (I) The CV of each attribute within each individual animal (box plot), and across animals (diamonds). The median is displayed as the middle line, the 25 % and 75 % quantiles as boxes, and the 10 % and 90 % quantiles as whiskers. The underlying source data can be accessed in the “Figure 2-source data” file.

One important reason for variability across animals is known from previous studies: the size of animals [42]. We therefore investigated the impact of animals’ size on the six behavioural locomotion attributes and found that both IS speed and IS distance were correlated with the body length of a larva, whereas other attributes were not affected (Fig. 2-E1). This finding is in line with previous work [42]. We therefore decided to correct for the body length in all following analyses by dividing any distance measured by the average body length of the individual larva in question. Distances are reported in body lengths (bl), speeds and velocities in body lengths per second (bl/s).

### 2.3 Bending and head-casting behaviour is more variable within individuals than peristaltic forward locomotion

After investigating the inter-individual variability, we explored the intra-individual variability in behaviour (Fig. 2G-I). We found that attributes related to the peristaltic forward movement such as the IS speed, IS distance and IS interval were much less variable than attributes associated with directional changes such as the absolute bending and HC angles (Fig. 2I; as the HC rate is calculated as a single value per individual it is excluded from this analysis). This finding suggests that peristaltic forward movement is a highly stable, regular behaviour with little variation, whereas directional changes are much more variable. Interestingly, the directional changes, but not the peristaltic forward movement, were much more variable within each individual than across individuals (Fig. 2I).

### 2.4 Attributes of peristaltic forward locomotion are stable over time within individuals

Next, we wondered how stable the behaviour of individual animals was over time. To this end, we determined each of the six behavioural attributes for each individual in the first and in the third minute of the three-minute video, and asked whether the behaviour in either minute would be correlated with that in the other. We detected mild correlations of the HC rate as well as the absolute bending and HC angles, and strong correlations of the IS speed, the IS distance and the IS interval (Fig. 3). This suggests that the behaviour of individual animals, though variable, was nevertheless remarkably consistent over time: the fastest animals in the first minute were also the fastest in the third minute, and the animals making the largest HCs in the first minute tended to do the same in the third minute. This finding may suggest that individual larvae have particular behavioural traits that are different between, but consistent within, individual animals.

**Fig. 3:**
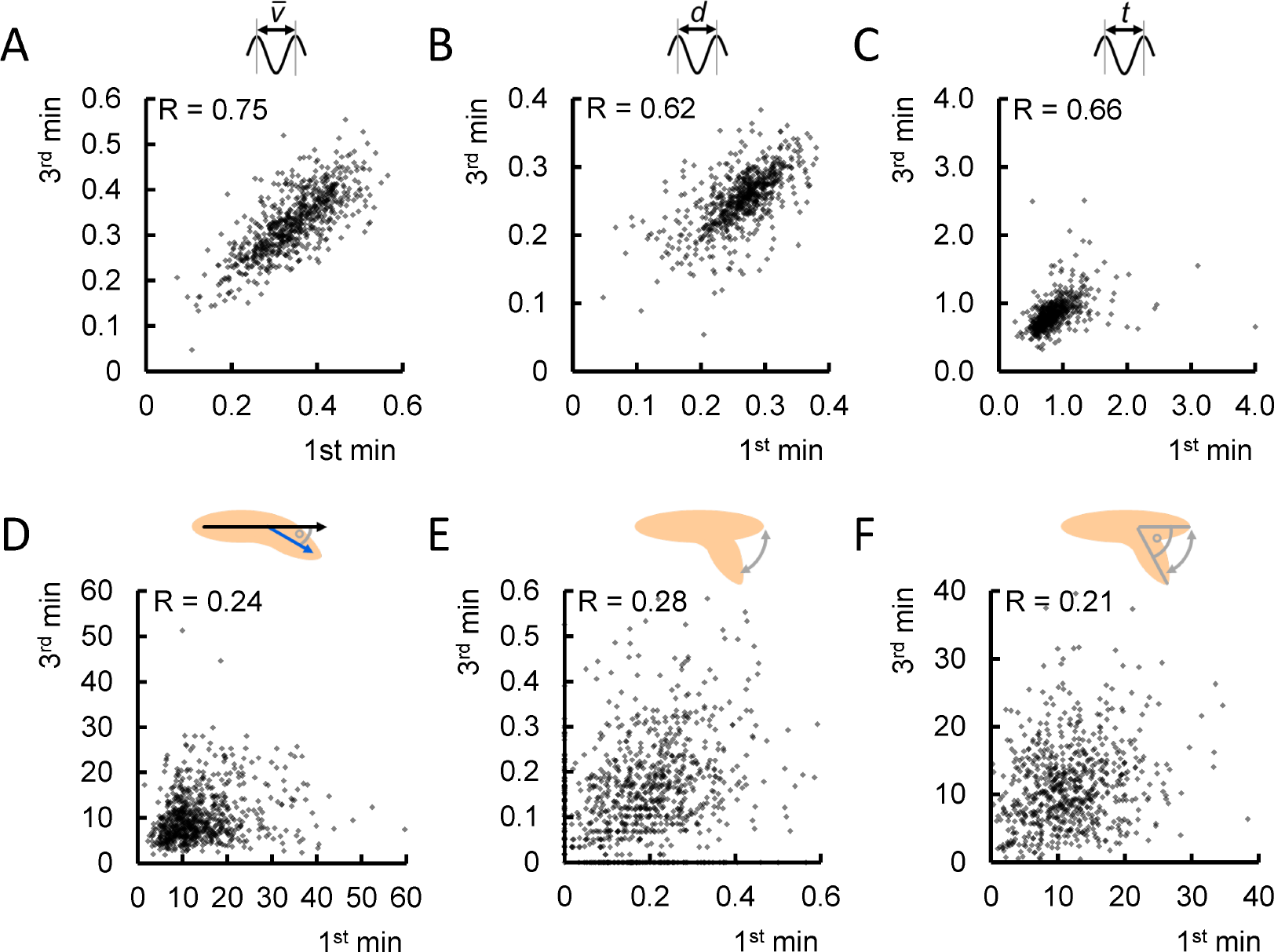
Basic locomotor attributes differ in how consistent they are over time. We determined each behavioural attribute for each individual independently for the first and the third minute of the video. We uncovered differences across behavioural attributes: we found moderate to strong correlations for (A) the IS speed, (B) the IS distance and (C) the IS interval, but only very weak, yet significant correlations for (D) the absolute bending angle, (E) the HC rate and (F) the absolute HC angle. Correlations are determined by SC tests. The underlying source data, as well as the results of the statistical tests, can be accessed in the “Figure 3-source data” file.

### 2.5 The Cirl mutation reduces speed, bending and head-casting behaviour

To put the newly developed technology to the test, we performed detailed analyses of larval locomotion and its modulations in response to genetic manipulation. To this end, we first chose the Latrophilin homolog Cirl, which is broadly expressed throughout the larval nervous system. Importantly, *Cirl*^*KO*^ mutant larvae are known to exhibit a strong locomotion deficit compared to wild type animals as well as a genomic *Cirl*^*Rescue*^ control [40], which has not, however, been studied in detail thus far.

We tracked and analysed the behaviour of 225 mutant and 147 rescue control larvae (Fig. 4). First, we confirmed the previously observed phenotype in crawling speed (Fig. 4B) [40]. Then, we analysed mutant and control animals regarding the six behavioural attributes described above. We found that the IS speed of the mutants was reduced compared to the control (Fig. 4C). Notably, a lower speed can come about in two ways: the larvae may make a slower peristaltic movement, resulting in a higher IS interval, or they may move less per peristaltic cycle, resulting in a lower IS distance. In the present case, we found only a small difference in the IS distance and a large difference in the IS interval (Fig. 4D-E). An even closer look into the run behaviour revealed that mutant animals are disturbed in their regular peristaltic cycle (see below).

**Fig. 4:**
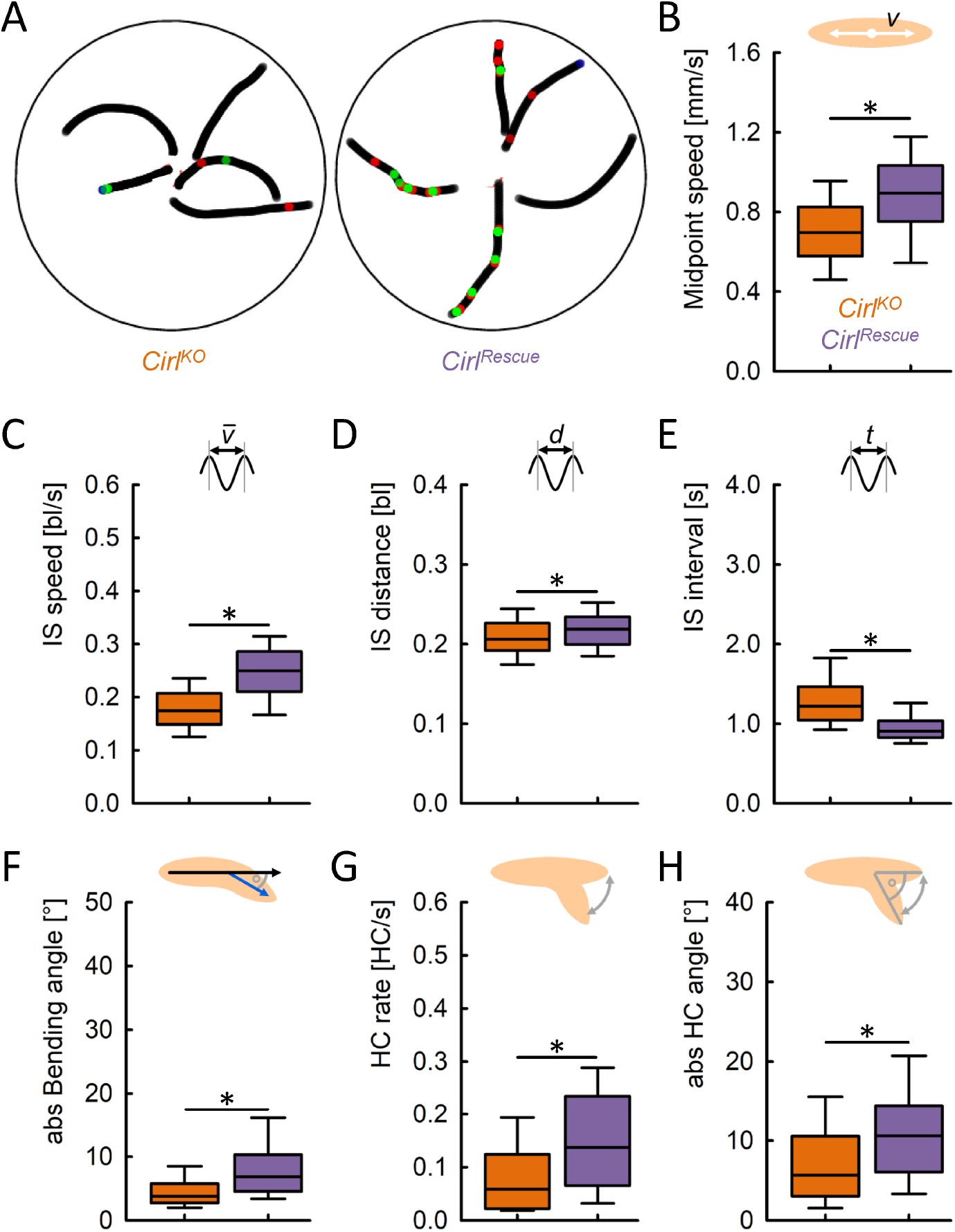
The Cirl mutation reduces speed, bending and head-casting behaviour. We analysed basic locomotor properties of the *Cirl*^*KO*^ mutant and a *Cirl*^*Rescue*^ control. (A) Representative sample tracks of five individual mutant and five control larvae. Green and red dots indicate left and right HCs, respectively. (B) The mutants displayed a strongly reduced midpoint speed compared to the control animals, confirming earlier results. (C) Also limited to run phases, the speed of mutants was reduced. (D) The IS distance was decreased, and (E) the IS interval increased. (F) In addition, the mutants were on average bending less than the control animals, which manifested itself as a decreased (G) HC rate and (H) absolute HC angle. The median is displayed as the middle line, the 25 % and 75 % quantiles as boxes, and the 10 % and 90 % quantiles as whiskers. Asterisks indicate significant differences in MW tests. The underlying source data, as well as the results of the statistical tests, can be accessed in the “Figure 4-source data” file.

In addition to the expected phenotype in speed, we detected a reduced absolute bending (Fig. 4F), which was accompanied by a reduced HC rate, as well as a reduced absolute HC angle, in the *Cirl*^*KO*^ mutant larvae (Fig. 4G-H). That is, the mutants made fewer and smaller HCs, and were bending less. This suggests that mutant larvae are generally impaired in the bending of the body and in directional changes, possibly due to impaired functioning of the chordotonal organs. This result was surprising, as in a previous study a qualitative description of the *Cirl*^*KO*^ mutant locomotion implied an increased number of HCs [40].

We also analysed the variability of each of the behavioural attributes within individuals, and observed increased variability in the IS speed and the IS interval in *Cirl*^*KO*^ mutant larvae compared to the controls (Fig. 4-E1).

### 2.6 A machine-learning approach reveals additional modulations of locomotion in *Cirl*^*KO*^ mutants

So far, our analysis had focused on the six basic locomotion attributes. But the *Cirl*^*KO*^ mutation may affect behaviour in further ways. We therefore extended our analysis to include additional behavioural attributes, including angular velocities of the head and tail, accelerations of head and tail, as well as the coefficients of variation whenever applicable. A full list of the 30 behavioural attributes used can be found in Tables 2-3 in the Materials and Methods section. We then employed an unbiased, machine-learning-based approach to determine the most decisive attributes that differentiate mutants and control larvae. Random forests work by building a large number of decision trees, each of them trying to classify each individual animal according to either one genotype or the other. Importantly, random forests are not only useful in classifying data, but also in estimating which attributes are most important for the classification. For more details, see the Materials and Methods section. Applying this approach to our data set, we were able to discriminate mutant and control larvae with an accuracy of 0.91 (Fig. 5A). Our assessment of which of the behavioural attributes were the most important in discriminating the behaviour of the two genotypes found that most of the top behavioural attributes were those already covered in our previous analysis: the IS speed, the IS interval, the absolute bending and HC angle, as well as the coefficient of variation of the IS speed and the IS interval (Fig. 5B). This result confirmed our analysis so far. However, the random forest method also revealed four additional behavioural attributes: the distance travelled by the larva, the absolute angular speeds of the head vector (HV) and tail vector (TV), as well as the coefficient of variation of the forward velocity. A closer look revealed that the *Cirl*^*KO*^ mutant larvae travelled shorter distances (Fig. 5C), consistent with their reduced speed. Furthermore, the mutants had decreased absolute HV and TV angular speeds (Fig. 5D-E), consistent with the general observation of decreased bending and turning rates in the mutant. Finally, we did not observe any clear difference in intra-individual variability in the tail forward velocity between mutant and rescue (Fig. 5F). The last result implies that the random forest can use attributes to differentiate individual animals that do not display obvious differences when plotted as box plots.

**Table 1:** Description of all basic body attributes used in this study.

| Name | Unit | Description | Random Forest |
| --- | --- | --- | --- |
| Spine points (x, y) | mm | The 12 spine points of the animal from tail to head. The first pair of coordinates is the position of the rear end and the 12th spine point indicates the front end of the animal. All spine point positions are convoluted by 0.3 seconds. | No |
| Contour points (x, y) | mm | The 24 contour points of the animal's body. All contour point positions are convoluted by 0.3 seconds. | No |
| Body length | mm | The sum of distances between all 12 spine points of the animal. | No |
| Head vector (x, y) |  | The vector between the 9<sup>th</sup> and the 11<sup>th</sup> spine point. | No |
| Tail vector (x, y) |  | The vector between the 2<sup>nd</sup> and the 6<sup>th</sup> spine point divided by its magnitude. | No |

**Table 2:** Description of all bending- and head-cast-related behavioural attributes used in this study.

| Name | Unit | Description | Random Forest |
| --- | --- | --- | --- |
| Bending angle | $^{\circ}$ | The angle between the larva's tail vector, and a vector from the middle of the larva to its head. Positive values indicate the larva bending to the left, negative values bending to the right. | Yes |
| Absolute bending angle | $^{\circ}$ | The absolute value of the bending angle. | Yes |
| CV absolute bending angle | | The coefficient of variation (CV) of the absolute bending angle, calculated as $100 * \frac{s(ABA)}{\overline{ABA}}$, where $s(ABA)$ is the standard deviation, and $\overline{ABA}$ is the mean of the absolute bending angle. | Yes |
| HV angular speed | $\frac{^{\circ}}{s}$ | The angular speed of the head vector (HV). Positive values indicate movement of the head to the right, negative values to the left. | Yes |
| Absolute HV angular speed | $\frac{^{\circ}}{s}$ | The absolute value for the HV angular speed. | Yes |
| CV absolute HV angular speed |  | The coefficient of variation (CV) of the absolute HV angular speed, calculated as exemplified above for the absolute bending angle. | Yes |
| HV angular acceleration | $\frac{^{\circ}}{s^2}$ | The angular acceleration of the head vector (HV). Positive values indicate movement of the head to the right, negative values to the left. | Yes |
| TV angular speed | $\frac{^{\circ}}{s}$ | The angular speed of the tail vector (TV). Positive values indicate movement of the tail to the right, negative values to the left. | Yes |
| Absolute TV angular speed | $\frac{^{\circ}}{s}$ | The absolute value for the TV angular speed. | Yes |
| CV absolute TV angular speed |  | The coefficient of variation (CV) of the absolute TV angular speed, calculated as exemplified above for the absolute bending angle. | Yes |
| TV angular acceleration | $\frac{^{\circ}}{s^2}$ | The angular acceleration of the tail vector (TV). Positive values indicate movement of the tail to the right, negative values to the left. | Yes |
| IS angle | $^{\circ}$ | The inter-step (IS) angle indicates how much the orientation of the animal changes within one cycle of peristaltic forward movement. It is defined as the angle between a larva's head vector at one step minus the head vector at the previous step. A step is defined as the local maximum of the tail forward velocity during runs. Positive values indicate that the larva changes its orientation to the left, negative values that it changes its orientation to the right. | Yes |
| Absolute IS angle | $^{\circ}$ | The absolute value of the IS angle. | Yes |
| CV absolute IS angle |  | The coefficient of variation (CV) of the absolute IS angle, calculated as exemplified above for the absolute bending angle. | Yes |
| HC rate | $\frac{HC}{s}$ | The number of head casts (HC) per second. | Yes |
| HC angle | $^{\circ}$ | The head-cast (HC) angle indicates how much the orientation of the animal changes upon each HC. It is defined as the difference between the bending angle at the end and the beginning of the HC. Positive values indicate that the larva changes its orientation to the left, negative values that it changes its orientation to the right. | Yes |
| Absolute HC angle | $^{\circ}$ | The absolute value of the HC angle. | Yes |
| CV absolute HC angle |  | The coefficient of variation (CV) of the absolute HC angle, calculated as exemplified above for the absolute bending angle. | Yes |

**Table 3:** Description of all speed-related attributes used in this study.

| Name | Unit | Description | Random Forest |
| --- | --- | --- | --- |
| Midpoint speed (mm) | $\frac{mm}{s}$ | The speed of the larva's midpoint, independent of direction. The midpoint is defined as the 6<sup>th</sup> spine point. | No |
| Midpoint speed (body length) | $\frac{bl}{s}$ | The speed of the larva's midpoint (measured in body lengths per second), independent of direction. | No |
| CV midpoint speed |  | The coefficient of variation (CV) of the midpoint speed, calculated as exemplified above for the absolute bending angle. | No |
| Head forward velocity | $\frac{bl}{s}$ | The velocity of the larva's head point in a forward direction, that is in the direction of the head vector. | Yes |
| CV head forward velocity |  | The coefficient of variation (CV) of the head forward velocity, calculated as exemplified above for the absolute bending angle. | Yes |
| Tail forward velocity | $\frac{bl}{s}$ | The velocity of the larva's tail point in a forward direction, that is in the direction of the tail vector. | Yes |
| CV tail forward velocity |  | The coefficient of variation (CV) of the tail forward velocity, calculated as exemplified above for the absolute bending angle. | Yes |
| IS distance (mm) | $mm$ | The inter-step (IS) distance measures the distance of the larva's midpoint between two steps. A step is defined as the local maximum of the tail forward velocity during runs. | No |
| IS distance (body length) | $bl$ | The inter-step (IS) distance, measured in body lengths per second. | Yes |
| CV IS distance |  | The coefficient of variation (CV) of the IS distance, calculated as exemplified above for the absolute bending angle. | Yes |
| IS interval | $s$ | The inter-step (IS) interval measures the time between two steps. A step is defined as the local maximum of the tail forward velocity during runs. | Yes |
| CV IS interval |  | The coefficient of variation (CV) of the IS interval, calculated as exemplified above for the absolute bending angle. | Yes |
| IS speed (mm) | $\frac{mm}{s}$ | The average midpoint speed between two steps. | No |
| IS speed (body length) | $\frac{bl}{s}$ | The average midpoint speed between two steps, measured in body length per second. | Yes |
| CV IS speed |  | The coefficient of variation (CV) of the IS speed, calculated as exemplified above for the absolute bending angle. | Yes |
| Distance travelled | $bl$ | The distance of the complete path a larva travels during the observation period. | Yes |
| Distance from startpoint | $bl$ | The maximal distance a larva reaches from its starting-point during the observation period. The starting-point is the location of the larva when it is first identified by the tracker. | Yes |

**Fig. 5:**
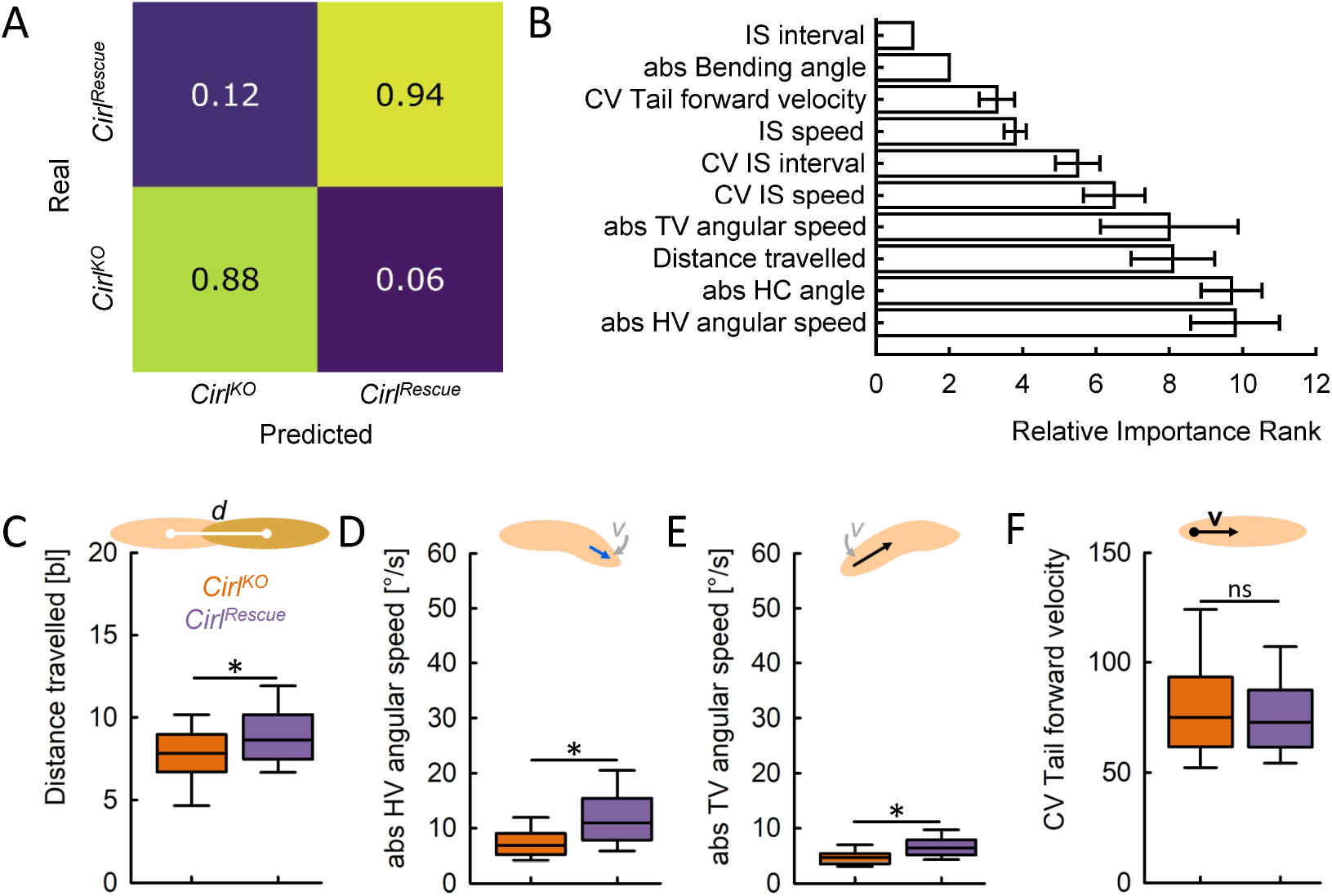
Discriminating *Cirl*^*KO*^ mutant and control animals animals by means of a random forest. We applied the random forest approach to discriminate *Cirl*^*KO*^ mutant animals from control animals. (A) Using a set of 30 behavioural attributes, the random forest was able to discriminate the two genotypes with an accuracy of 0.91. Displayed is the confusion matrix showing predicted versus actual affiliations of individual larvae to the two genotypes. (B) The top ten most important behavioural attributes for discriminating the two genotypes with the random forest, sorted by the average importance rank across 10 repetitions of the random forest approach. (C-F) Four additional behavioural attributes found by the random forest were (C) the distance travelled by the larvae, (D) the angular speed of the head vector (HV), (E) the angular speed of the tail vector (TV), as well as (F) the coefficient of variation (CV) of the velocity of the tail in a forward direction. Displayed are the mean and the 95 % confidence interval in (B), and the median as the middle line, the 25 % and 75 % quantiles as boxes, and the 10 % and 90 % quantiles as whiskers in (C-F). Asterisks indicate significant differences in MW tests. The underlying source data, as well as the results of the statistical tests, can be accessed in the “Figure 5-source data” file.

To confirm that these top ten behavioural attributes were indeed the critical ones for describing the behaviour of *Cirl*^*KO*^ mutants and control animals, we ran the random forest algorithm again, but this time with only those ten attributes. The algorithm was able to discriminate the two genotypes with an accuracy of 0.91, just as high as with the full set of behavioural attributes. The ten attributes in question were thus shown to be fully sufficient to discriminate between the genotypes. In turn, when we ran the random forest algorithm with the remaining 20 attributes and excluding the top ten, the accuracy was still 0.83. This suggests some redundancy in our analysis, with multiple attributes providing related information such that the algorithm can employ substitute attributes to discriminate between the genotypes even when the most decisive attributes are excluded.

### 2.7 *Cirl*^*KO*^ mutant larvae have an altered rhythmic peristaltic cycle

In order to understand the origin of the *Cirl*^*KO*^ phenotype in crawling speed, we analysed the temporal behavioural patterns of individual animals (Fig. 6). Just like wild-type larvae (Fig. 1D), *Cirl*^*Rescue*^ controls displayed a very regular rhythm in their forward velocity, indicated by a smooth, wave-like curve with local maxima of very similar heights that we detected as steps, and always one local minimum between two steps (Fig. 6A, top, Rich media file 1). *Cirl*^*KO*^ mutants, in contrast, regularly exhibited a pattern of two local minima with one very small local maximum between two steps (Fig. 6A, bottom, Rich media file 2), resulting in IS intervals that were longer and more variable than those of *Cirl*^*Rescue*^ controls (Fig. 4E, 4-E1C). For further analyses, we defined each period between two steps with only one local minimum in the tail forward velocity as normal stepping, and each period with two local minima as ‘stumble’ stepping. A quantification revealed that stumble stepping was much more common in mutant than in control larvae, and that their proportion increased with increased IS intervals (Fig. 6B-C).

**Fig. 6:**
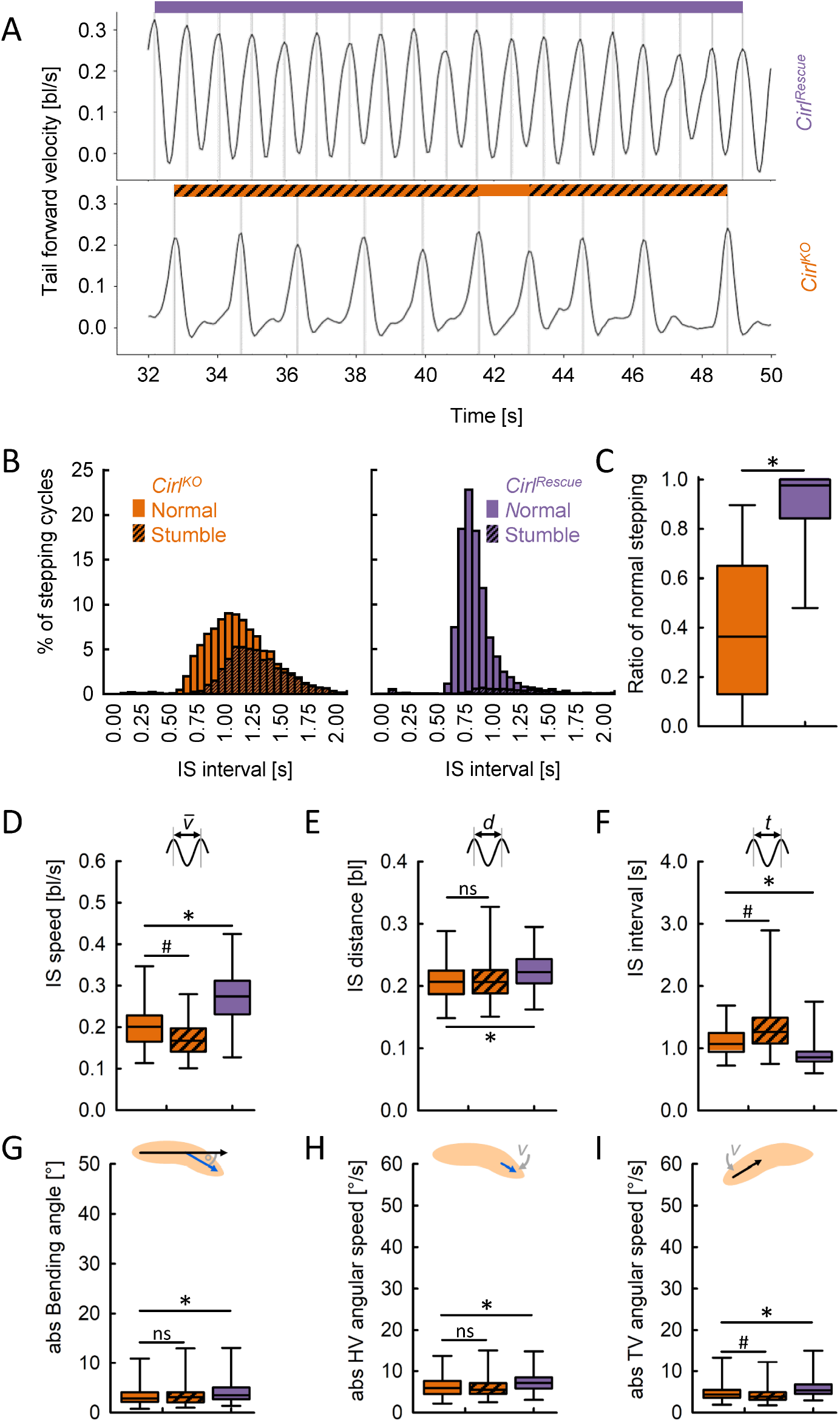
*Cirl*^*KO*^ mutant larvae have an altered rhythmic peristaltic cycle. (A) The forward velocity of the tail in each one sample *Cirl*^*Rescue*^ (top) and *Cirl*^*KO*^ larva (bottom). Periods of normal stepping are indicated by a coloured stripe on top of the curve, periods of stumble stepping by a hatched stripe. (B) Distribution of the IS interval between the normal and stumble stepping cycles for each genotype. (C) *Cirl*^*KO*^ individuals have a much lower ratio of normal stepping cycles (and thus a higher ratio of stumble stepping) than *Cirl*^*Rescue*^ individuals. (D-I) Within-animal comparison of various behavioural attributes of mutant animals during normal or stumble stepping, and between-animal comparison between mutant and control animals during normal stepping. Periods of stumble stepping in control animals were omitted due to their low number. (D) IS speed, (E) IS distance, (F) IS interval, (G) absolute Bending angle, (H) absolute head vector (HV) angular speed, (I) absolute tail vector (TV) angular speed. Displayed are the median as the middle line, the 25 % and 75 % quantiles as boxes, and the 10 % and 90 % quantiles as whiskers. Asterisks indicate significant MW tests, hash symbols significant WS tests. The underlying source data, as well as the results of the statistical tests, can be accessed in the “Figure 6-source data” file.

Next, we directly compared the animals’ behaviour specifically during normal stepping and stumble stepping. We found that larvae were slower and had longer IS intervals, but unchanged IS distances during stumble stepping compared to normal stepping (Fig. 6D-F). In other words, stumble steps took longer to complete, but were not larger. Likewise, neither the animals’ bending nor the angular speed of their head vector was different between normal and stumble steps; only the angular speed of the tail vector was slightly decreased (Fig. 6G-I). This suggests that the phenotype in bending and turning that we observed was largely independent of the irregular stepping behaviour of the *Cirl*^*KO*^ mutants.

In summary, we found that the *Cirl*^*KO*^ mutants’ phenotype in crawling speed can be at least partially attributed to an unusual stepping pattern that causes a slower peristaltic forward locomotion, whereas the phenotype in bending and turning is largely independent.

### 2.8 Optogenetic activation of dopamine neurons triggers stopping and bending

Next, we investigated how the acute optogenetic activation of sets of neurons affected the locomotion of individual larvae. First, we applied our unbiased approach to the behaviour of *Drosophila* larvae upon optogenetic activation of dopamine neurons. To this end, we used larvae that expressed the highly sensitive effector Channelrhodopsin2-XXL (ChR2-XXL) [43] in a broad subset of dopaminergic neurons, as defined by the *TH-Gal4* driver strain (experimental genotype). Groups of larvae were allowed to move freely over a Petri dish for 30 s in darkness, followed by 30 s of blue light in order to trigger the activation of the dopaminergic neurons, and another 60 s of darkness. The behaviour of the animals was compared to that of heterozygous driver and effector controls that did not express ChR2-XXL (Fig. 7). Upon light stimulation, animals of all three genotypes increased their absolute bending, suggesting that this was a response to the sudden light. However, the experimental genotype did so to a much greater extent than the genetic controls, suggesting that this stronger reaction was caused by the activation of the dopaminergic neurons (Fig. 7B, Rich media file 3).

**Fig. 7:**
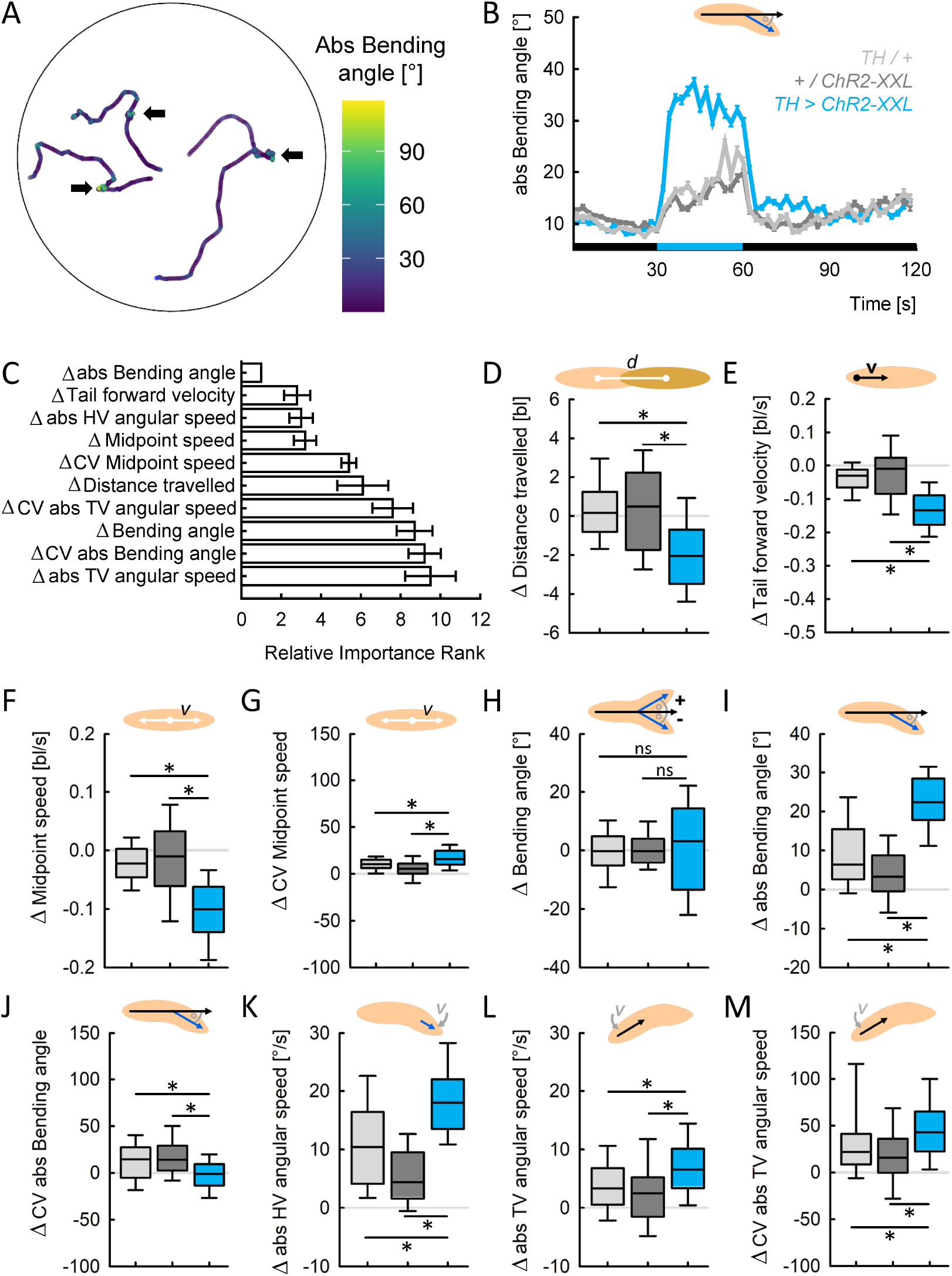
Optogenetic activation of dopamine neurons triggers stopping and bending. Larvae of the experimental genotype (*TH >ChR2-XXL*), the driver control (*TH / +*) and the effector control (*+ / ChR2-XXL*) were recorded during free locomotion for 30 s in darkness, followed by 30 s of light stimulation and 60 s of darkness. (A) Sample tracks of three individuals, colour-coded according to the absolute bending angle. Phases of stopping and increased bending are indicated by brighter colour (arrows). (B) The absolute bending angle over time. Black and blue stripes on the X-axis indicate periods of darkness and blue light stimulation, respectively. (C) 30 behavioural attributes during light stimulation were normalized to each individual’s behaviour before light activation (called Δ-values). Shown are the top ten most important Δ-values for discriminating the three genotypes by means of random forest, sorted by the average importance rank across 10 repetitions. The top ten Δ-values provided by random forest were (D) the distance each larva travelled, (E) the velocity of the tail in a forward direction, (F) the speed of the midpoint, as well as (G) its coefficient of variation (CV), (H) the bending angle, (I) the absolute bending angle, as well as (J) its CV, (K) the absolute angular speed of the head vector (HV), (L) the absolute angular speed of the tail vector (TV), as well as (M) its CV. Displayed are the mean and the 95 % confidence interval in (B-C), and the median as the middle line, the 25 % and 75 % quantiles as boxes, and the 10 % and 90 % quantiles as whiskers in (D-M). Asterisks indicate significant differences in MW tests. The underlying source data, as well as the results of the statistical tests, can be accessed in the “Figure 7-source data” file.

In order to evaluate the reaction of the animals to the light stimulation and the light-induced neuronal activation, we normalized each individual animal’s behaviour during the 30 s of light stimulation by subtracting its behaviour in the 30 s of darkness before. The normalized values of 30 behavioural attributes (called Δ-values) were fed into the random forest. The random forest classified the three genotypes with an accuracy of 0.90 and provided the top ten most important attributes that differed between the three genotypes in their change in response to the light stimulation (Fig. 7C). In detail, we found a more strongly reduced distance travelled by larvae of the experimental genotype than of the genetic controls and accordingly a more strongly reduced tail forward velocity and midpoint speed (Fig. 7D-F). Interestingly, the intra-individual variability in the midpoint speed, as determined by the coefficient of variation, was increased in all three genotypes, but more in the experimental genotype (Fig. 7G). In addition, the random forest approach provided three behavioural attributes regarding the animals’ bending. For the bending angle, which is positive if a larva is bent to the left and negative if bent to the right, we found no significant differences, but a strongly increased scatter of the data in the experimental genotype (Fig. 7H). Fittingly, the absolute bending angle, not taking the direction into account, was increased in all three genotypes, but much more so in the experimental genotype (Fig. 7I). The intra-individual variability in the absolute bending angle was increased only in the genetic controls, but not in the experimental genotype (Fig. 7J). Finally, the angular speeds of the animals’ head and tail vectors also differed between the genotypes: both angular speeds were increased more strongly in the experimental genotype than in the controls (Fig. 7K-L). Regarding the tail vector angular speed, the intra-individual variability was also more strongly increased in the experimental genotype (Fig. 7M).

In summary, activating a broad set of dopaminergic neurons caused the animals to slow down markedly and to bend and rotate their bodies (for a time-resolved view of the three genotypes’ behaviour, see Fig. R7-E1).

### 2.9 Individuals reduce speed consistently upon repeated activation of dopamine neurons

In wild-type animals we observed that behavioural attributes associated with peristaltic forward crawling were more consistent within each animal than attributes associated with bending and HC behaviour (Fig. 2-3). We therefore wondered whether this would also be true for the dopamine-induced changes in behaviour that we had observed. To tackle this question, we repeated the previous experiment with the experimental genotype expressing ChR2-XXL in *TH-Gal4* neurons, but this time we activated the neurons three times for 10 s, each time followed by 60 s of darkness (Fig. 8). Then we determined the changes in the four behavioural attributes that had previously been found to be most important by the random forest method (Fig. 7C): tail forward velocity, midpoint speed, absolute bending angle, and absolute angular speed of the head vector. On average, the animals reacted very similarly to the three neuronal activations (Fig. 8). As regards the changes in behaviour within each animal, however, we found differences between the behavioural attributes. We observed mild to moderate correlations across the three neuronal activations for the tail forward velocity and the midpoint speed (Fig. 8B, D; Fig. 8-E1B, D), but no correlations for the absolute bending angle and the absolute angular speed of the head vector (Fig. 8F, G; Fig. 8-E1F, G).

**Fig. 8:**
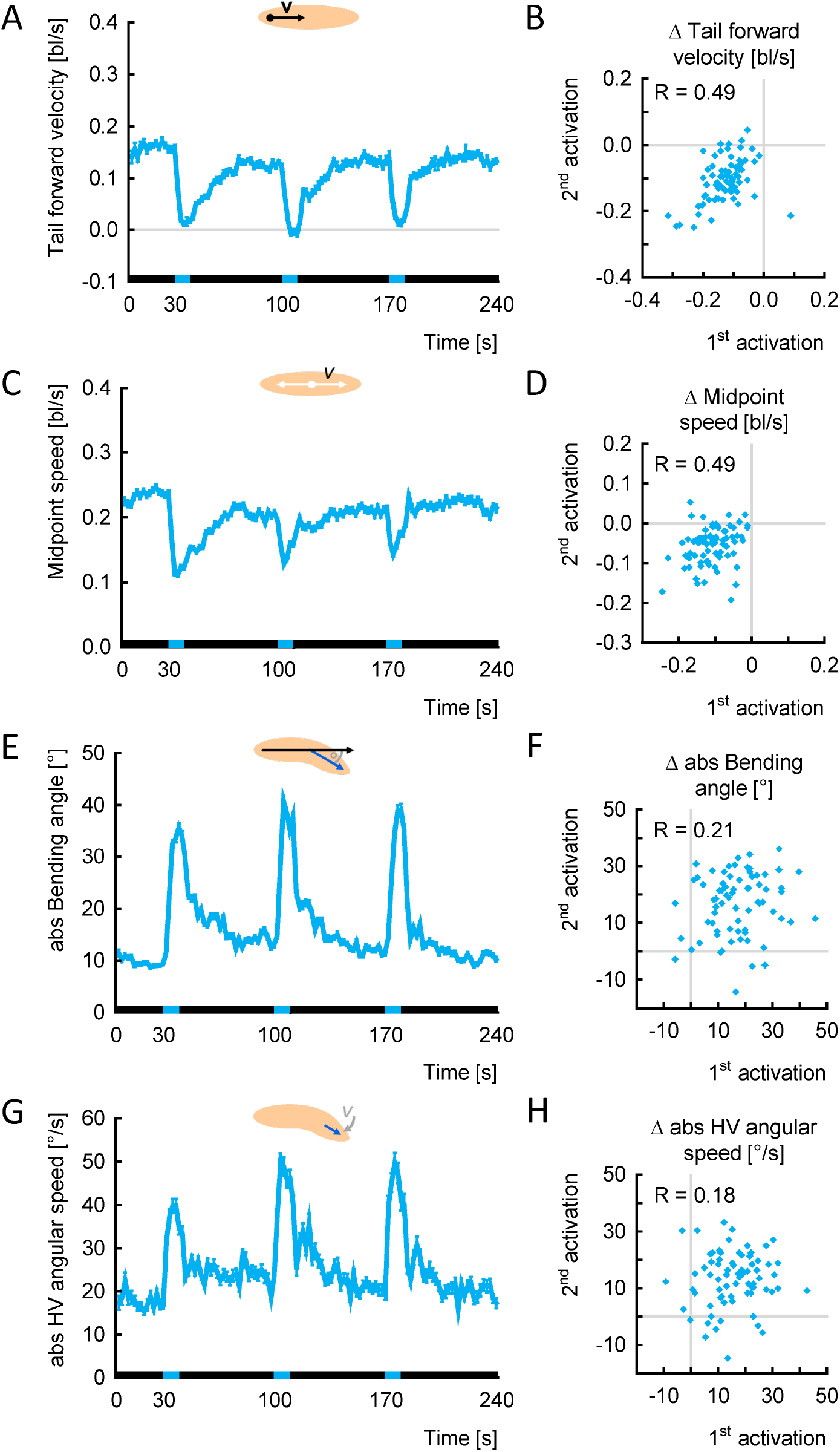
Individuals reduce speed consistently upon repeated optogenetic activation of dopamine neurons. Larvae of the experimental genotype (*TH >ChR2-XXL*) were recorded during free locomotion for 30 s in darkness, followed by three cycles of 10 s of light stimulation and 60 s of darkness. (A) The tail forward velocity over time. Black and blue stripes on the X-axis indicate periods of darkness and blue light stimulation, respectively. (B) The tail forward velocity during light stimulation was normalized to each individual’s behaviour in the 10 s before light activation (called Δ-values). The Δ-velocities of the first and second activation were positively correlated. (C-D) As in (A-B), but for the midpoint speed. (E-F) As in (A-B), but for the absolute bending angle. (G-H) As in (A-B), but for the HV absolute angular speed. Displayed are the mean and the 95 % confidence interval in (A,C,E,G), and the individual Δ-values in (B,D,F,H). Correlations were determined by SC tests. The underlying source data, as well as the results of the statistical tests, can be accessed in the “Figure 8-source data” file.

Thus, we indeed found that the reduction in speed upon activation of dopamine neurons was consistent within individual animals, whereas the increases in bending and lateral head movement were not. This further stresses the distinction between forward crawling and bending behaviour with respect to intra-individual variability.

### 2.10 Optogenetic activation of ‘mooncrawler’ neurons triggers backward crawling

In order to probe the utility of our approach further, we applied it to an analogous experiment to that described above, but activating a very different set of neurons. This time we chose the driver strain *R53F07-Gal4*, which reportedly covers a rather broad set of interneurons in the ventral nerve cord (VNC), the insects’ functional analog of the spinal cord, including the so-called moonwalker descending neurons (MDNs) [44]. Optogenetically activating these neurons (henceforth called ‘mooncrawler’ neurons) had caused larvae to crawl backwards in a previous study [44], but a in-depth analysis of their behaviour is lacking. Therefore, we crossed *R53F07-Gal4* to the *UAS-ChR2-XXL* effector strain, performed the same experiment as described above, and compared the animals’ behaviour with the respective genetic controls (Fig. 9). Light stimulation did indeed cause backward crawling, as indicated by a negative forward velocity, in animals of the experimental genotype but not of the genetic controls (Fig. 9B, Rich media file 4).

**Fig. 9:**
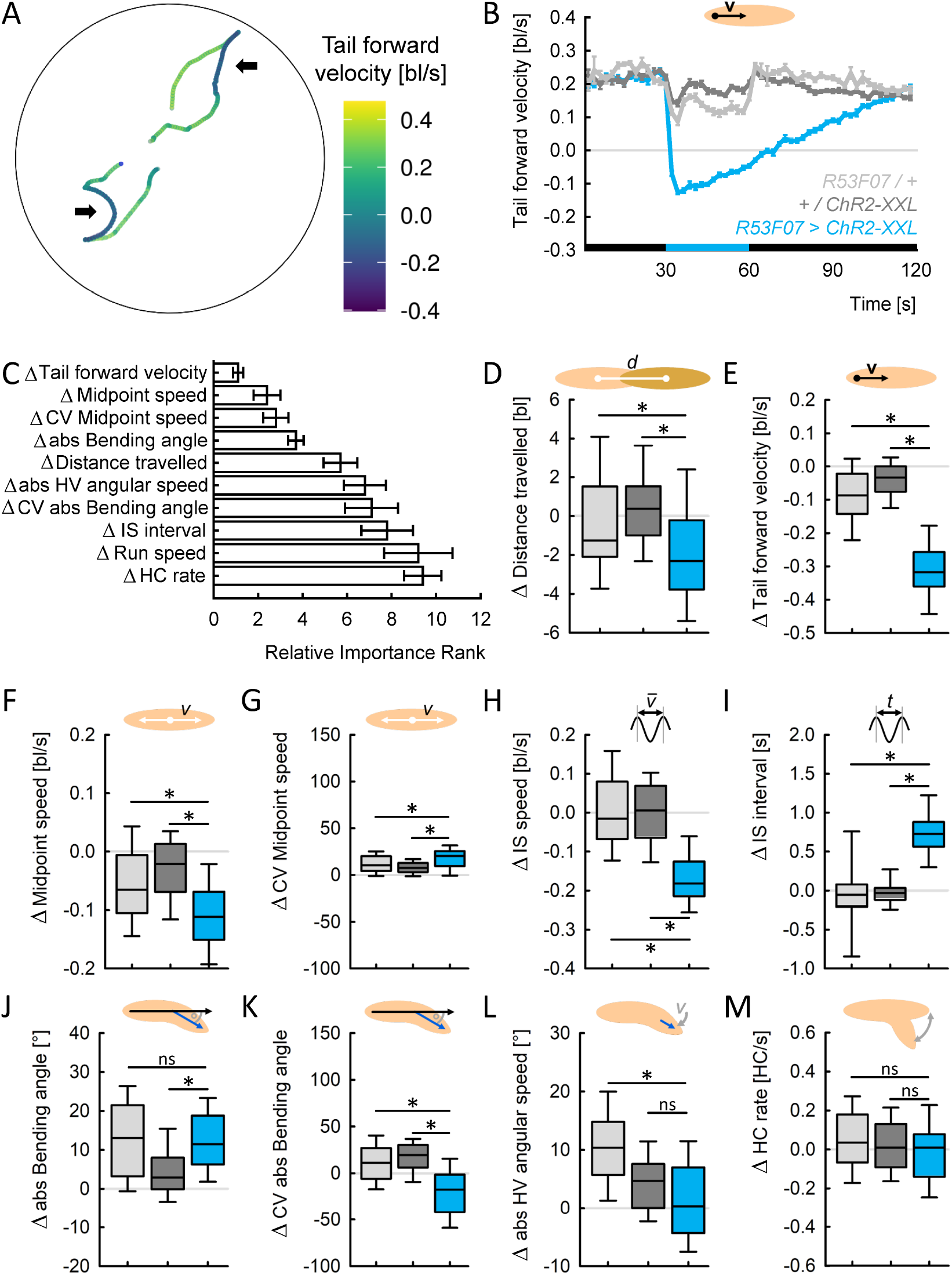
Optogenetic activation of ‘mooncrawler’ neurons triggers backward crawling. Larvae of the experimental genotype (*R53F07 >ChR2-XXL*), the driver control (*R53F07 / +*) and the effector control (*+ / ChR2-XXL*) were recorded during free locomotion for 30 s in darkness, followed by 30 s of light stimulation and 60 s of darkness. (A) Sample tracks of two larvae, colour-coded according to their tail forward velocity. Phases of backward locomotion are indicated by darker colour (arrows).(B) The tail forward velocity over time. Black and blue stripes on the X-axis indicate periods of darkness and blue light stimulation, respectively. (C) 30 behavioural attributes during light stimulation were normalized to the individual’s behaviour before light activation (called Δ-values). Shown are the top ten most important Δ-values. The top ten Δ-values were (D) the distance each larva travelled, (E) the velocity of the tail in a forward direction, (F) the speed of the midpoint, as well as (G) its coefficient of variation (CV), (H) the IS speed, (I) the IS interval, (J) the absolute bending angle, as well as (K) its CV, (L) the absolute angular speed of the head vector (HV), and (M) the HC rate. Displayed are the mean and the 95 % confidence interval in (B-C), and the median as the middle line, the 25 % and 75 % quantiles as boxes, and the 10 % and 90 % quantiles as whiskers in (D-M). Asterisks indicate significant differences in MW tests. The underlying source data, as well as the results of the statistical tests, can be accessed in the “Figure 9-source data” file.

We applied the same strategy as before, fed the Δ-values of behavioural attributes into the random forest (accuracy 0.83), and obtained the top ten attributes that differed between the three genotypes in their change in response to the light stimulation (Fig. 9C). Similarly to the case of dopaminergic neurons, we found the distance travelled, the tail forward velocity and the midpoint speed to be more strongly decreased in the experimental genotype than in the genetic controls, and the intra-individual variability of the midpoint speed to be more strongly increased (Fig. 9D-G). In contrast to the previous experiment, however, the IS speed and the IS interval were also among the top ten attributes: we found the IS speed to be strongly reduced, and the IS interval to be strongly increased, in the experimental genotype but not the genetic controls (Fig. 9H-I). Again, we found the animals’ bending to be important, yet in a different way from the previous experiment: the absolute bending angle was reported as an important attribute by random forest, but the experimental genotype was different only from the effector control, not from the driver control (Fig. 9J). Thus, there was no evidence that the activation of the *R53F07-Gal4* neurons were responsible for the change in the absolute bending angle. Interestingly, the intra-individual variability in the absolute bending angle was increased in the genetic controls but decreased in the experimental genotype (Fig. 9K). Although the change in the average bending was not significantly different from the genetic controls, therefore, activating the *R53F07-Gal4* nneurons made the bending less variable. Finally, the angular speed of the head vector as well as the HC rate were reported by the random forest. However, for the angular speed we found a significant difference in the experimental genotype only in relation to the driver, but not the effector control (Fig. 9L), and no significant differences in the HC rate at all (Fig. 9M).

### 2.11 Individuals differ in the timing of their switching gears from backward to forward locomotion

A closer look at the temporal patterns of the behavioural attributes induced by the two sets of neurons activated revealed striking differences. The activation of dopaminergic neurons triggered a number of behavioural changes that quickly reverted back to normal as soon the light stimulation ended (Fig. 7-E1). In contrast, activating the *R53F07-Gal4* neurons triggered behavioural changes of very different, more complex dynamics (Fig. 9-E1). For example, the tail forward velocity dropped quickly to the negative upon light onset and started to become more positive gradually over the course of 90 s (Fig. 9-E1A). The midpoint speed and IS speed, in contrast, continued to drop slowly until a minimum was reached about 30 to 40 s after light onset (Fig. 9-E1B-C). The absolute bending angle was increased in all three genotypes during light stimulation. But whereas it reverted back to normal upon light offset in the genetic controls, it remained high for another 30 s in the experimental genotype before gradually decreasing (Fig. 9-E1E). What is the cause of these complex temporal patterns?

We compared the temporal pattern of the tail forward velocity across individual animals (Fig. 10-E1). Whereas all the individuals switched to backward crawling within a few seconds after light onset, the reverse switch to forward crawling was highly variable, with some animals not reverting at all within the recording time (Fig. 10A). In other words, most larvae continued to crawl backwards even after the optogenetic activation of the *R53F07-Gal4* neurons had finished, and each reverted to forward crawling in its own time. On average, those animals that did revert to forward locomotion within the recording time crawled backwards for about 51 s (Fig. 10B). This also means that our previous analysis was confounded by mixing unknown numbers of forward- and backward-crawling individuals at any given time point. Therefore, we determined the switching points in each individual animal (Fig. 10-E1) and compared the individual animals’ behaviour while the animals were crawling backward (i.e. between the switching points) with their behaviour while crawling forward (Fig. 10C).

**Fig. 10:**
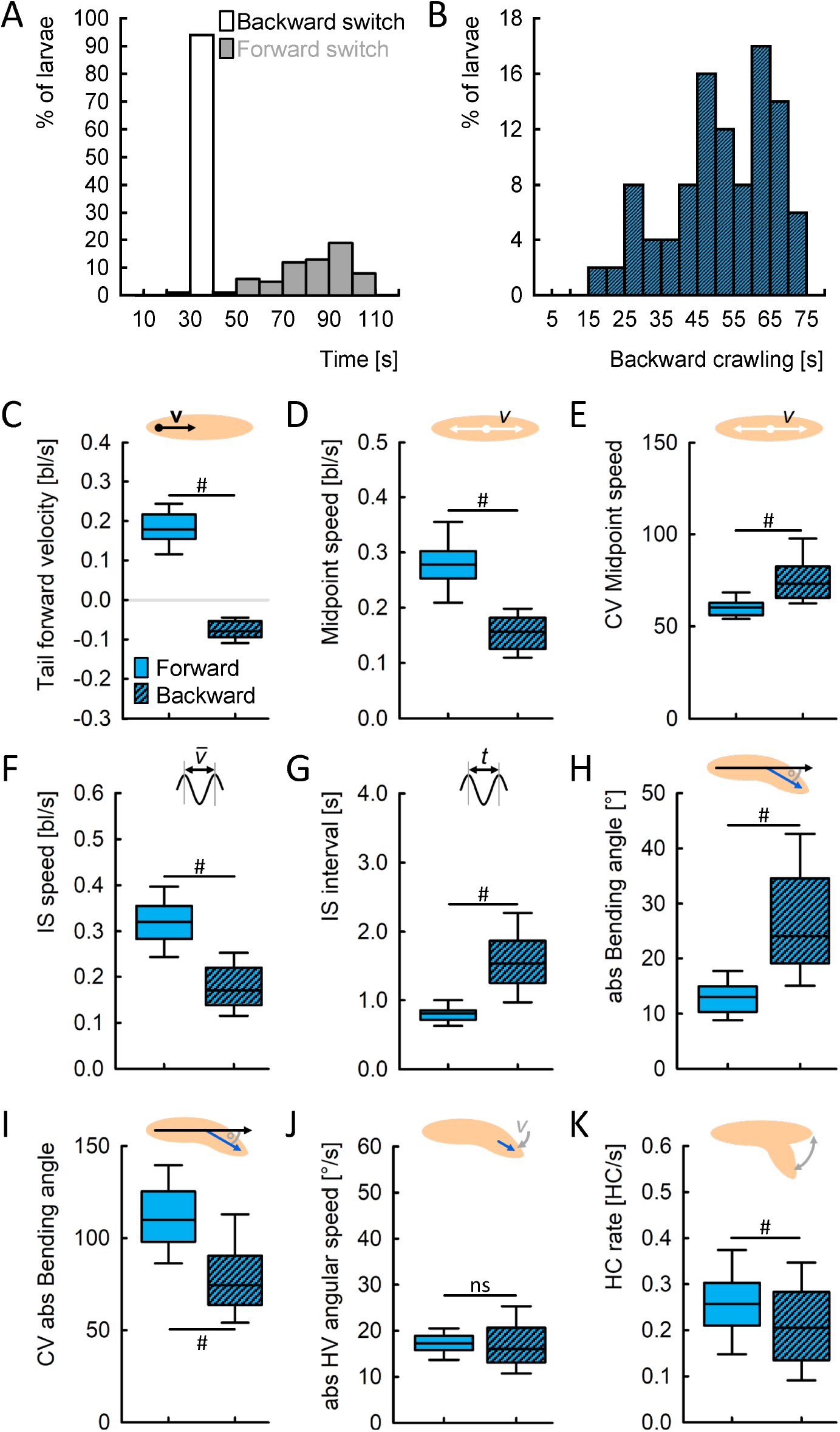
Backward locomotion is characterized by reduced yet more variable speed, and increased yet less variable bending. (A) Distribution of the switching points to backwards crawling (white) and forward crawling (grey). N = 97, 64. (B) Distribution of backward crawling durations, only taking those animals into account that perform both switches, i.e. that start to crawl backward and switch back to forward crawling later (N = 58). (C-K) Within-animal comparison of the backward crawling phase (i.e. between the switching points) and the forward crawling phases, regarding (C) the tail forward velocity, (D) the midpoint speed, (E) its coefficient of variation (CV), (F) the IS speed, (G) the IS interval, (H) the absolute bending angle, (I) its CV, (J) the HV absolute angular speed, and (K) the HC rate. Displayed are the median as the middle line, the 25 % and 75 % quantiles as boxes, and the 10 % and 90 % quantiles as whiskers. Hash symbols indicate significant differences in WS tests. The underlying source data, as well as the results of the statistical tests, can be accessed in the “Figure 10-source data” file.

We found that the midpoint speed was reduced, but its intra-individual variability was increased, while animals crawled backwards (Fig. 10D-E). Taking only run phases into account, the IS speed was reduced and the IS interval was accordingly increased, during backward crawling (Fig. 10F-G). The absolute bending angle was increased, with reduced intra-individual variability, while larvae crawled backwards (Fig. 10H-I). Finally, we found no differences between forward and backward crawling regarding the angular speed of the head vector, but a reduced HC rate during backward crawling (Fig. 10J-K).

In summary, backward crawling was characterized by longer IS intervals, leading to a reduced and more variable speed, and by increased but less variable bending of the larvae, compared to forward crawling.

### 2.12 Switches between forward and backward locomotion are characterized by changes in speed, and peaks in bending and head-casting behaviour

Finally, we wondered how specifically the animals’ behaviour was altered during the switches between forward and backward locomotion. To this end, we aligned the data of all the animals either to the switch to backward crawling, or to the switch to forward crawling (Fig. 11A). This alignment revealed that both midpoint speed and IS speed rapidly dropped at the switch to backward crawling, followed by a slow further decrease in the following seconds. Both speeds decreased until a few seconds before the switch to forward crawling and quickly rose again after the switch (Fig. 11B-C). Fittingly, the IS interval increased after the backward switch and decreased again after the forward switch (Fig. 11D). The absolute bending angle rapidly increased precisely at the backward switch and stayed constant afterwards. At the forward switch, the absolute bending angle peaked and then dropped quickly back to normal levels (Fig. 11E). The absolute angular speed of the head vector did not change at all at the backward switch but peaked at the forward switch (Fig. 11F). Finally, the HC rate peaked at both switching points, albeit on different absolute levels (Fig. 11G).

**Fig. 11:**
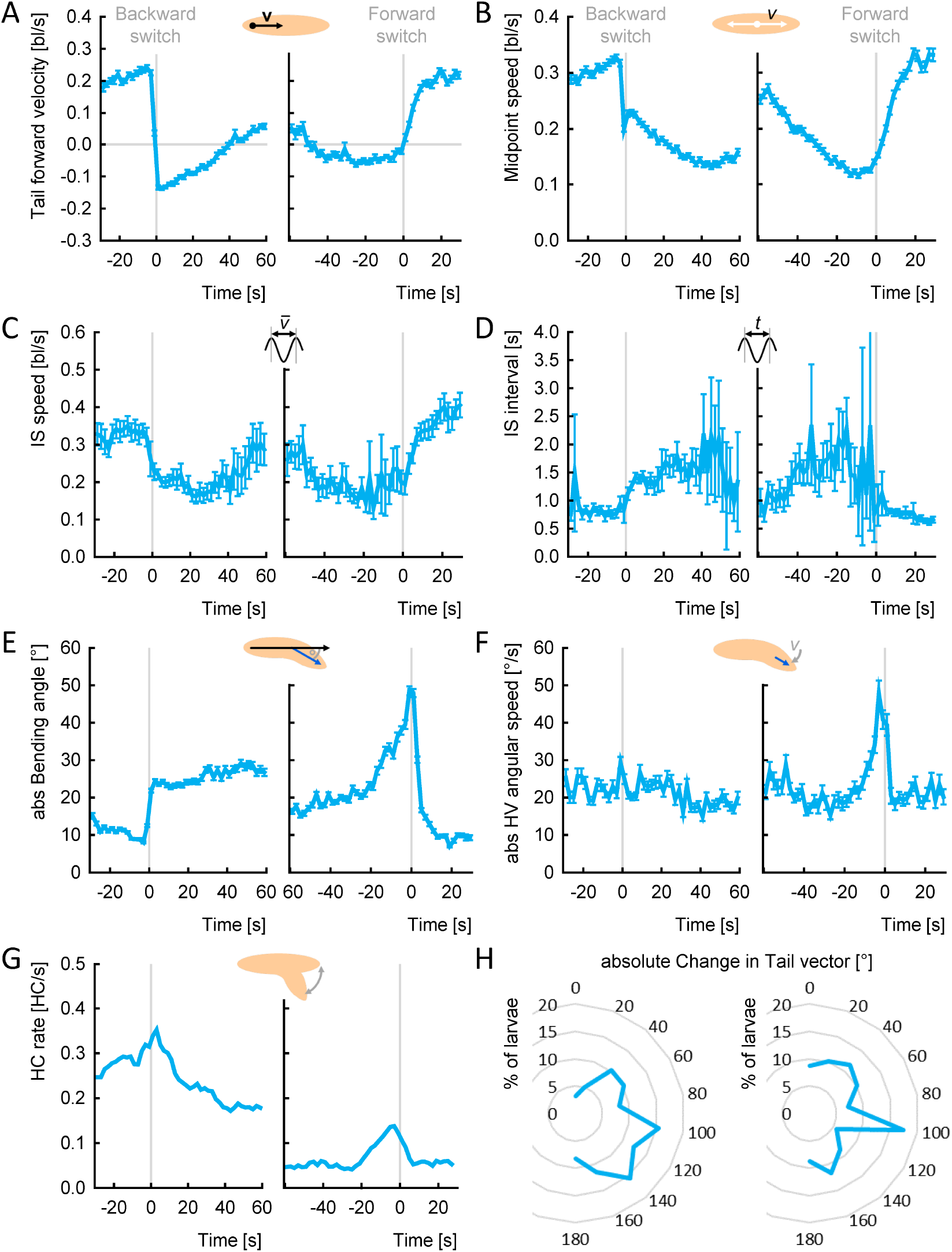
Switching between forward and backward locomotion (and *vice versa*) is characterized by changes in speed and peaks in bending and head-casting. (A-G) The tracks of animals from the experimental genotype (*R53F07 >ChR2-XXL*) were aligned either to the switch from forward to backward crawling (left, “Backward switch”), or to the switch from backward to forward crawling (right, “Forward switch”). Only animals that performed the respective switch were considered (N = 97, 64). (A) Tail forward velocity, (B) midpoint speed, (C) IS speed, (D) IS interval, (E) absolute bending angle, (F) absolute head vector (HV) angular speed, (G) HC rate. Displayed are the mean and 95 % confidence intervals of all data within each 2 s bin, except for the HC rate which was calculated within each 2 s bin by dividing all frames with a HC by all frames in the bin. (H) The orientation of the animal’s body, indicated by the tail vector, was compared in the 10 s before and after a switch. Displayed is the percentage of animals that changed their orientation by the respective absolute angle, in brackets of 20°. The underlying source data can be accessed in the “Figure 11-source data” file.

In order to determine how the direction of crawling changed at each switching point, we calculated the change in the tail vector in the 10 s before and after each switch (Fig. 11H). We found that in both cases the larvae changed their crawling direction by about 100 °, suggesting that either switch was followed by a movement approximately orthogonal to the previous direction.

### 2.13 Individuals change speed consistently upon repeated backward, but not forward switches

We characterized backward switches by a decrease in speed and an increase in absolute bending angle, and forward switches by an increase in speed, and peaks in absolute bending angle and absolute angular speed of the head vector. How stable were these changes for repeated switches across and within individuals? To address this question, we stimulated the *R53F07-Gal4* neurons three times for 10 s and aligned all animals either to the backward or to the forward switch (Fig. 12). We found that the average behaviour of the animals was astonishingly consistent across all three backward and forward switches. Looking at individual behaviour with regard to the changes in tail forward velocity and midpoint speed, we observed mild to moderate correlations across repeated backward switches (Fig. 12A, C; Fig. 12-E1A, C), but not across repeated forward switches (Fig. 12B, D; Fig. 12-E1B, D). A notable exception was a significant moderate correlation in the midpoint speed between the second and third forward switch (Fig. 12-E1D). In contrast, no correlations were found for the absolute bending angle and the absolute angular speed of the head vector (Fig. 12E-H; Fig. 12-E1E-H). The IS speed, IS interval and HC rate were not included in this analysis, as steps and HCs occur too rarely within the short observation periods used for this analysis.

**Fig. 12:**
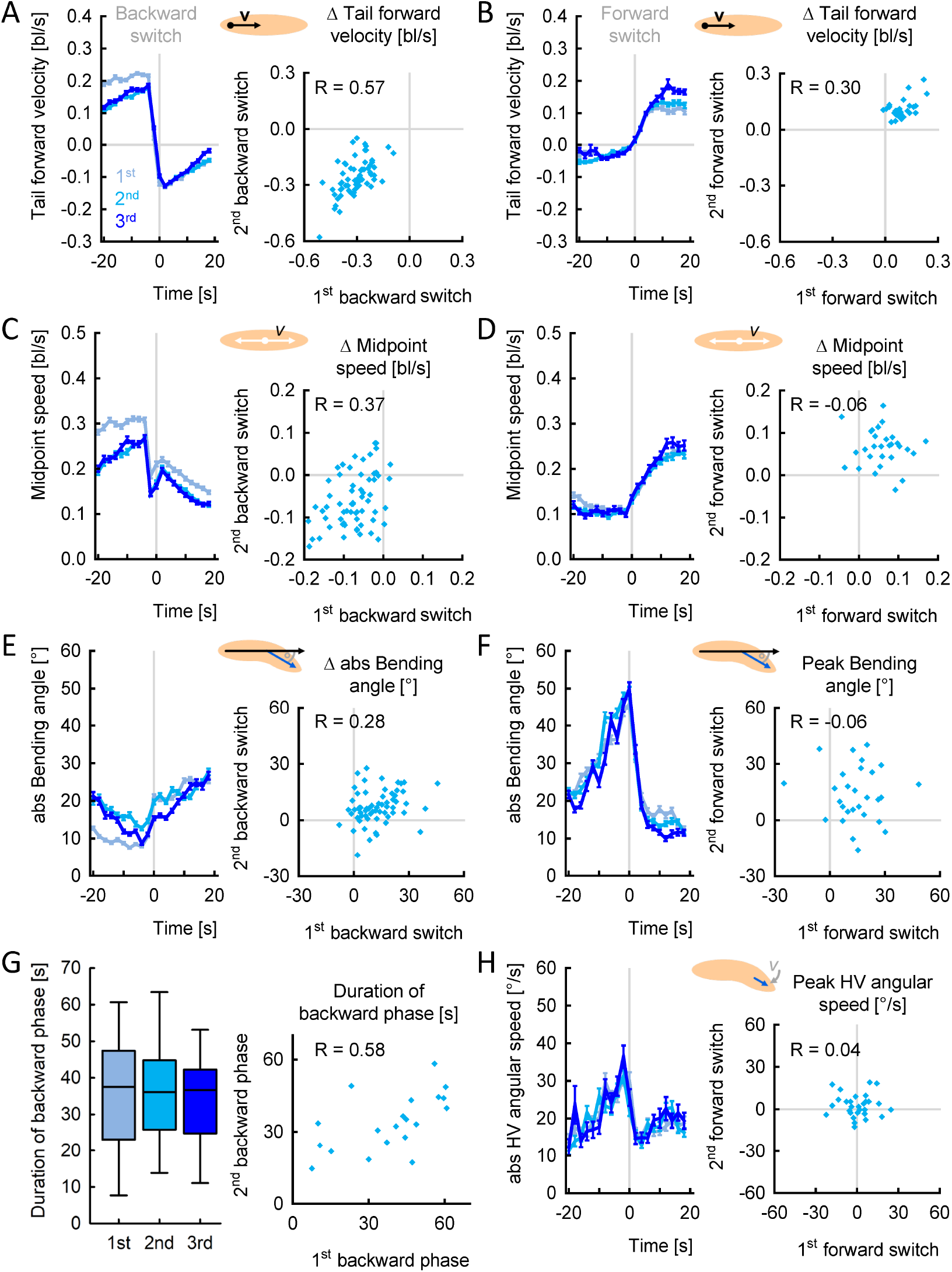
Individuals change speed consistently upon repeated backward, but not forward switches. Larvae of the experimental genotype (*R53F07 >ChR2-XXL*) were recorded during free locomotion for 30 s in darkness, followed by three cycles of 10 s of light stimulation and 60 s of darkness. Individual data were aligned either to the switch from forward to backward crawling (A,C,E), or to the switch from backward to forward crawling (B,D,F,H). (A) (left) The tail forward velocity over time, aligned to the first, second and third backward switch. (right) The tail forward velocity during the 10 s after the switch was normalized to each individual’s behaviour in the 10 s before the switch (called Δ-values). The Δ-velocity of the first and second activation was positively correlated. (B) As in (A), but for the forward switches. (C-D) As in (A-B), but for the midpoint speed. (E) As in (A), but for the absolute bending angle. (F) To quantify the peak of the absolute bending angle at forward switches, we normalized the behaviour in the 10 s around the switch with the 5 s before and the 5 s after (called peak values). (G) (left) The duration of the backward crawling phases following the first, second and third neuronal activation. (right) The durations of the first and second backward phases were positively correlated. (H) As in (F), but for the HV absolute angular speed. As no particular change in the angular speed is seen upon backward switches, we quantify only the forward switches. Displayed are the mean and the 95 % confidence interval in (A,C,E,G), and the individual Δ-values, or peak-values, in (B,D,F,H). Correlations were determined by SC tests. The underlying source data, as well as the results of the statistical tests, can be accessed in the “Figure 12-source data” file.

In addition, we determined the duration of each of the three backward crawling phases upon each activation of *R53F07-Gal4* neurons in individual animals, and found positive correlations that were significant between the first and the second, as well as between the second and third backward phases (Fig. 12G; Fig 12-E1G). Between the first and third backward phase too, a positive correlation was observed; this remained non-significant due to the sample size of only seven animals that completed both backward phases.

In summary, we again found that changes in speed upon neuronal activation were more consistent within individuals than changes in bending or lateral head movement, just as we found to be the case with respect to dopamine neurons.

## 3 Discussion

In this study, we demonstrate the utility of the IMBA with four examples: our findings provide new insights into variability in locomotor behaviour, the complex behavioural phenotypes of an adhesion GPCR mutation, the effects of repeatedly activating dopamine neurons on locomotion, and the ‘switching of gears’ between forward and backward locomotion. In the following paragraphs, we first discuss the implications of our findings and the possible future directions for each of these cases, before summarizing the more general potential of the IMBA.

### 3.1 Bending and head-casting behaviour is variable; peristaltic locomotion is stable

Our analysis revealed that peristaltic forward locomotion is characterized by very low intra-individual variability and high consistency over time (Fig. 2-3). Bending and head-casting behaviour, in contrast, is much more variable, and less stable. We note that, while similar findings have led to the suggestion that peristaltic forward locomotion would thus be best suitable for high-throughput phenotyping [42], we found reliable effects of transgenic manipulations on bending and HC behaviour too (Fig. 4, 7, 9). In any event, when we repeatedly activated the same set of neurons in the same individual animals, we found mild yet significant correlations for these repetitions for peristaltic forward locomotion, but not for bending and head-casting behaviour. This was true for both dopamine neurons and mooncrawler neurons (as covered by the *R53F07-Gal4* driver) (Fig. 8, 12). This suggests that individual larvae not only have a fairly stable, inherent rhythm of peristaltic locomotion, but they also modify this rhythm in a relatively idiosyncratic manner upon repeated neuronal activation. Bending and head-casting behaviour, on the other hand, appears highly variable and is modulated in an unpredictable manner upon repeated neuronal activation.

The reason for this apparent difference in variability between peristaltic locomotion on the one hand and bending and head-casting behaviour on the other hand is not known. Interestingly, however, previous studies have found that peristaltic locomotion originates in both the anterior and posterior segments of the VNC, whereas head-casting behaviour and bending originate more anteriorly [45, 46]. In both cases, input from the brain is reportedly not required. Input from the brain is required, in contrast, for organizing peristalsis, bending and head-casting behaviour in a goal-directed manner [45]. Consistently with this, in previous studies we have demonstrated that “higher” brain functions, such as associative memories, almost exclusively modulate head-casting behaviour, but not peristaltic runs [27, 41, 47]. Overall, it therefore seems that the characteristics of peristaltic runs that are particularly relevant for speed are relatively stable features of a given individual animal, “hard-wired” in the VNC, whereas bending and head-casting behaviour, and thus changes in direction, are more flexible features that can be adaptively modulated by external stimuli and internal goals, modulations that apparently take place via the brain. We are curious whether such a distinction between speed-related, hard-wired, stable motor outputs from the VNC and direction-related, flexible, brain-modulated behaviours may turn out to be a general principle of behavioural control.

We note that intra-individual variability itself can be affected by specific experimental manipulations: for example, we found that the *Cirl*^*KO*^ mutants show an increased variability in their IS interval (Fig. 4-E1).The optogenetic activation of select sets of neurons can also elicit changes in the variability of behaviour (Fig. 7, 9). One might therefore expect there to be cases in which experimental manipulation specifically affects the variability of a behavioural measure, but not its average. This would be interesting in particular in relation to processes for which behavioural variability is of particular value, e.g. for generating unpredictable actions during escape, for generating novel actions in pursuit of desired outcomes in operant learning, or for adapting to changed environmental contingencies during reversal learning.

### 3.2 *Cirl*^*KO*^ mutants exhibit ‘stumble’ steps

We discovered a new, complex behavioural phenotype of Latrophilin/Cirl mutants. Cirl constitutes an aGPCR (adhesion G-protein-coupled receptor) expressed in larval chordotonal organs and mechanosensory organs crucial for proprioception [40]. Previously, it was known only that mutants were slower, as was confirmed by our analysis (Fig. 4). Here we found that the reason for the reduced speed was the abnormal ‘stumble’ stepping of the mutant animals, in which the peristaltic cycle was interrupted, and the time required to complete one cycle was therefore prolonged (Fig. 6). Interestingly, this different way of stepping was in itself rather regular, such that some mutant individuals always performed stumble steps (Fig. 6A).

The stumble stepping explained parts of the observed behavioural phenotype: the IS interval was increased, and the IS speed was thus decreased during stumble stepping compared to ‘normal’ stepping in the same individual mutant animals (Fig. 6D, F). However, the normal stepping phases of mutant and control animals also differed in these behavioural attributes, suggesting that the peristaltic cycles that we classified as ‘normal’ were partially impaired as well. Moreover, we found that mutant animals bend and turn less than control animals, a phenotype that was completely independent of the kind of stepping the animals performed (Fig. 6G-H).

The molecular mechanisms through which aGPCRs exert their function remain incompletely understood. However, it is known that chordotonal neuron-located Cirl suppresses cAMP levels to shape larval proprioception and thus behaviour [40, 48]. If and how Cirl has an impact in other cells that are part of the neuronal circuits that modulate locomotion remains elusive. Our analysis suggests that the correct functioning of the receptor is required for a normal, uninterrupted peristaltic cycle, but that even without the receptor larvae can eventually crawl forward. This is in line with published accounts showing decreased but existent proprioceptive capability [40]. It will be interesting to study animals that express genetically modified Cirl, cell-specific knock-outs or knock-downs to elucidate the Cirl-expressing cells that contribute to larval locomotion, and the ability of *cirl* mutants successfully to reach goals such as odours or tastants despite their impairment in crawling.

### 3.3 Activation of dopamine neurons makes larvae stop and turn

Across the animal kingdom, the dopaminergic system is closely associated with locomotion and the generation of movement [49, 50]. In humans, the progressive death of dopaminergic neurons is widely accepted as a cause of Parkinson’s disease, which results in difficulties in initiating movement [51–53]. Based on studies mostly of rodents, locomotion has been found to be controlled by interactions of several dopamine receptors in the ventral striatum, with some of them increasing, others inhibiting forward locomotion [54]. Turning behaviour can be induced by the unilateral administration of dopamine receptor agonists [55]. The administration of cocaine, which strongly activates the dopaminergic system, causes complex, concentration-dependent modulations of locomotion, including increased forward locomotion upon administration of lower concentrations of cocaine, and circling behaviour as well as a reduced forward movement upon repeated, or highly concentrated cocaine administration [56–58].

In *Drosophila* too, the dopaminergic system is linked to locomotion. Genetic fly models for Parkinson’s disease show similar impairments in locomotion, as well as the death of dopaminergic neurons as found in mammals [59]. Cocaine administration leads successively to a reduction in forward movement, circling behaviour and rotating, before the animals stop moving at all, similar to the case in mammals [60]. In a pioneering optogenetic study, activation of the dopamine neurons covered by *TH-Gal4* resulted in increased locomotion in flies that were little active before, and decreased locomotion in flies that were highly active [61]. Recently, a study has found the activity of specific dopamine neurons to be correlated with speed and turning of flies, suggesting that dopamine neurons both report and change specific movements [62].

Much less is known about the impact of dopamine on locomotion in *Drosophila* larvae. In this study, we optogenetically activated most dopamine neurons, and found that larvae stop forward locomotion and start strong bending (Fig. 7). Such behaviour would correspond to the behaviour of adult flies and mice after cocaine administration [56–58, 60], and to what was observed in highly active flies upon activation of dopamine neurons [61]. Our results are also in line with one of the very few studies that link dopamine function with locomotion in larvae: a transgenic knockdown of the dopamine receptor Dop1R1, one of the two *Drosophila* homologs of the vertebrate D1-type dopamine receptors, was reported to increase locomotion in larvae [63]. Our experimental treatment, in contrast, acutely increased dopamine signalling, leading to reduced forward locomotion.

Interestingly, the effects of the Dop1R1-knockdown were also observed when it occurred locally only in the mushroom body, one of the central brain regions in insects [63]. The dopamine neurons innervating the mushroom body play a crucial role in signalling reward or punishment information in associative learning [29, 64–68]. The mushroom body-innervating neurons covered by *TH-Gal4* are mostly linked to punishment signalling, with the reward-signalling dopamine neurons notably missing in the expression pattern [64, 69, 70]. This raises the question of what the activation of these neurons ‘means’ to the animals. On the one hand, stopping and turning can be interpreted as an avoidance behaviour in response to a sudden punishment signal (for a similar interpretation regarding the activation of dopamine-downstream neurons, see [71]). On the other hand, an excessive release of dopamine, as probably happens in our experiment, may cause malfunctions in the motor control pathways, similar to what was observed upon cocaine administration in adults [60]. The fine genetic tools available make the *Drosophila* larva an ideal study case to answer this question. Dopamine neurons can be manipulated in small subsets and even one by one [65–67] to find out which neuron causes which behavioural reaction.

### 3.4 Switching from forward to backward locomotion, and back again

In free, undisturbed locomotion, *Drosophila* larvae normally crawl forward, and backward crawling is very rarely observed. They do, however, perform short bouts of backward locomotion on encountering an obstacle or receiving a noxious stimulus to their head [72, 73]. Thus, backward locomotion can be seen as part of an avoidance reaction. In recent years, several Gal4 driver strains have been described that cover neurons which induce backward locomotion upon activation [44, 73, 74]. One of them, *R53F07-Gal4*, we used in the present study [44]. We described many facets of the behaviour of individual animals over extended phases of backward locomotion, including the bending and head-casting behaviour that had not been looked at in previous studies [44, 73, 74]. We found, for example, that during backward locomotion bending was increased in amplitude, but decreased in variability, and that the HC rate was decreased compared to forward locomotion (Fig. 10H-K). Moreover, we found that larvae did not revert to forward locomotion immediately after the optogenetic activation terminated. Rather, they continued to crawl backward for various time periods, and then quickly switched to forward locomotion (Fig. 10-E1). Both kinds of switches were accompanied by characteristic changes in speed and bending behaviours (Fig. 11, 13).

**Fig. 13:**
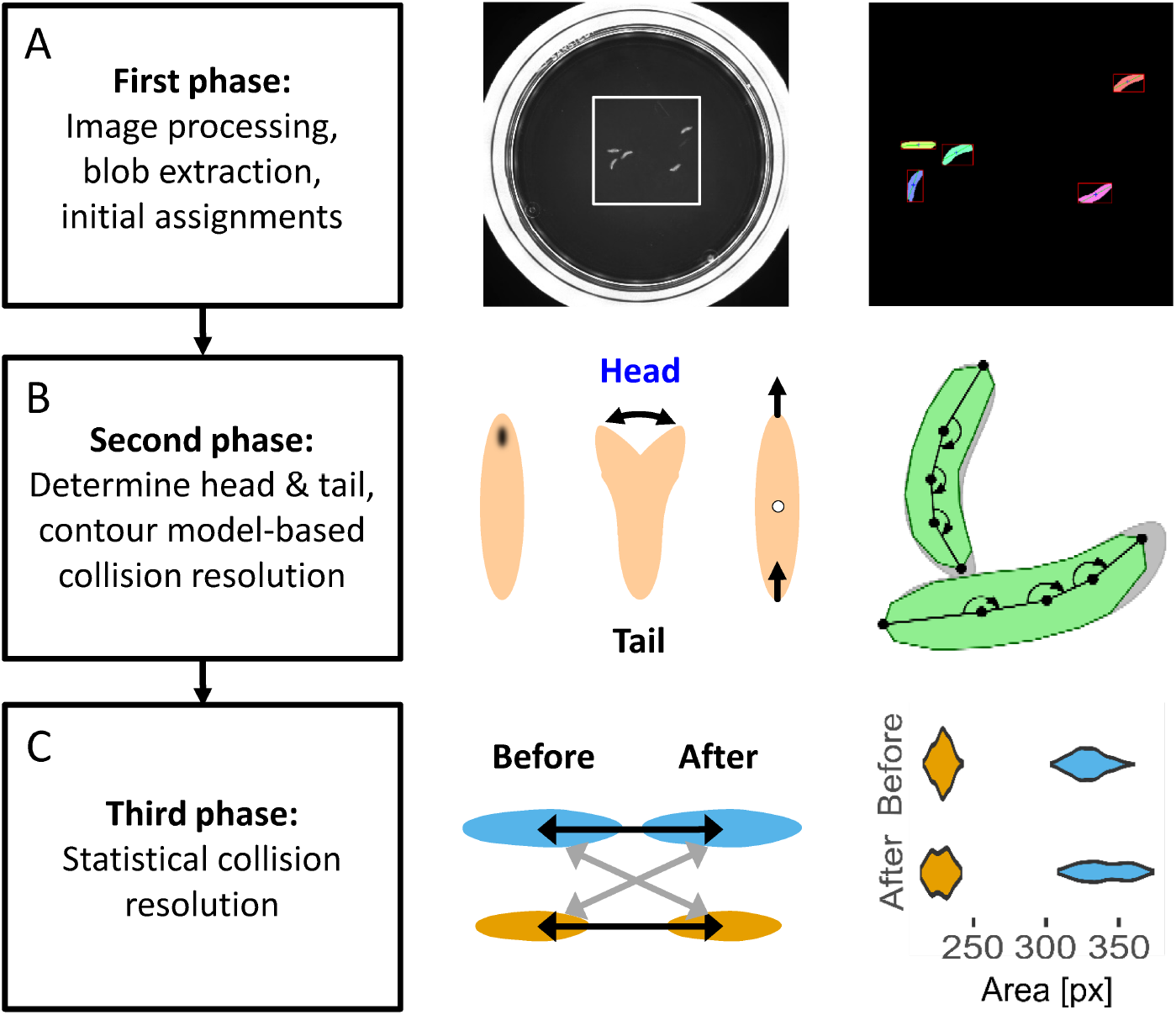
Overview of the tracking process. The IMBAtracker works through a sequence of three phases. (A) In the first phase, the image is processed: e.g. by background subtraction, objects (“blobs”) are detected by thresholding, and initial assignments of individual larvae are made before collision resolution). (B) In the second phase, the head and tail of each larva is determined, and a contour-based model of colliding larvae is applied to undertake a first round of collision resolution. (C) For collisions that could not be resolved in the second phase, the third phase adds a statistical collision-resolution approach that compares attributes such as the size of the larvae to match individuals before and after the collision.

On the neuronal level, both forward and backward locomotion are characterized by waves of activity in the motor neurons and interneurons of the segmented VNC, which travel from anterior to posterior during forward locomotion, and the other way around during backward locomotion [8, 45, 75, 76]. More specifically, two pairs of command-like neurons have been identified in the rather broad expression pattern of *R53F07-Gal4*, called mooncrawler descending neurons (MDNs), which inhibit forward and induce backward locomotion [74]. It is likely that the effects of activating *R53F07-Gal4* neurons come about through these MDNs. Two main effects of activating the MDNs have been proposed: on the one hand, one of their main downstream partners, called Pair1, stops forward locomotion by inhibiting the forward-movement-inducing A27h neuron; on the other hand, another downstream partner, A18b, induces backward locomotion specifically in the most anterior segment of the VNC [11, 14, 74]. Remarkably, both the MDNs and Pair1 neurons persist through metamorphosis, and induce analogous behavioural effects in adult flies [74, 77, 78].

It is a straight-forward assumption that the switches between forward and backward locomotion that we describe here may be caused by the backward-movement-inducing A18b and the forward-movement-inducing A27h neurons, respectively. The observation that larvae did not revert to forward locomotion as soon as the optogenetic activation ended suggests that activating the MDN→A18b pathway leads to backward locomotion that continues until forward locomotion is triggered again, potentially by A27h. Our behavioural approach, together with the available tools for fine circuit analysis, is suited to test this hypothesis.

### 3.5 Utility for analysing individual behaviour

The IMBA software introduced here can serve to track, visualize and analyse the behaviour of individual *Drosophila* larvae within groups. The IMBA is compatible with well-established, standard Petri-dish assays and requires no specific additional hardware beyond stable light conditions and a camera. Indeed, the present study features data from setups located both at the Leibniz Institute for Neurobiology and the University of Leipzig, which use different means of lighting, cameras and video quality. We also successfully analysed videos from other researchers with yet different conditions (not shown). Such robustness makes the IMBA a useful tool for studying a broad range of research questions using derivatives of this standard assay. In the present study, we focused on locomotion and used the IMBA to phenotype a mutant (Fig. 4-6) and study the effects of optogenetic neuronal activation (Fig. 7-12). Other future applications may include the screening of pharmacological effectors [79] and libraries for RNAi knockdown, and comparisons across inbred strains of Drosophila or across species.

A critical property of the IMBA that allows for such broad use is that we included a large number of behavioural attributes that can be adapted and extended by the user (the full list is available online under https://github.com/XXX). Among them are 22 attributes describing behaviour relative to a local stimulus. This makes the IMBA useful for experiments on behaviour towards stimuli such as odours, tastants, light or temperature, as well as for analysing stimulus-related behaviour after various types of learning experiment. Importantly, we combine this large number of behavioural attributes with an approach to selecting the most relevant ones for each particular data set in an unbiased way. Random forest analysis provides a ranking of behavioural attributes according to their importance for differentiating animals in the particular experimental conditions of the particular data set under study (compare Fig. 7C and 9C). This includes attributes that do not differ in their centres of distribution (and thus result in non-significant standard statistical tests) but rather in the distribution of the data (Fig. 5F, 7H).

The IMBA is versatile for analysing relatively fine-grained temporal patterns of behaviour. This has led us to discover stumble stepping in *Cirl*^*KO*^ mutants (Fig. 6). Indeed, the IMBA allows for the analysis of even more extreme kinds of ‘silly walks’ that it was not designed to study, such as the backward locomotion elicited by activation of MDN neurons (Fig. 9-12). Furthermore, the IMBA lends itself to protocols that involve repeated experimental stimulation, as in the case of the repeated activation of select sets of neurons (Fig. 8, 12). This should make it possible to establish larval versions of paradigms such as prepulse inhibition or repetition priming, which are workhorse paradigms in the experimental neuropsychology of rodents and humans. Last but not least, the IMBA allows data to be aligned to changes in behaviour (that is, to the time of action initiation rather than stimulus application), revealing effects that would otherwise be masked by differences in action-timing between individuals (Fig. 11, 12).

In summary, the IMBA is an easy-to-use toolbox providing an unprecedentedly rich view of behaviour and behavioural variability across the multitude of biomedical research contexts in which larval *Drosophila* can be used.

## 4 Material and Methods

### 4.1 Data sets

We used six different data sets for our analysis:

- A set of 48 videos with 875 individual animals entering the analysis. Wildtype larvae were tested for innate odour preference on Petri dishes of 15 cm over 3 minutes. These data were previously published in [27], and are re-analysed in this study regarding individual animals.
- A set of 126 videos with 372 individual animals entering the analysis. *Cirl*^*KO*^ and *Cirl*^*Rescue*^ larvae were tested for locomotion over 1 minute. For details, see below.
- A set of 50 videos with 253 individual animals entering the analysis. Larvae expressing ChR2-XXL in *TH-Gal4* neurons, together with the respective genetic controls, were tested for locomotion upon optogenetic activation over 2 minutes in total. For details, see below.
- A set of 56 videos with 257 individual animals entering the analysis. Larvae expressing ChR2-XXL in *R53F07-Gal4* neurons, together with the respective genetic controls, were tested for locomotion upon optogenetic activation over 2 minutes in total. For details, see below.
- A set of 22 videos with 122 individual animals entering the analysis. Larvae expressing ChR2-XXL in *TH-Gal4* neurons were tested for locomotion upon repeated optogenetic activation over 4 minutes in total. For details, see below.
- A set of 20 videos with 115 individual animals entering the analysis. Larvae expressing ChR2-XXL in *R53F07-Gal4* neurons were tested for locomotion upon repeated optogenetic activation over 4 minutes in total. For details, see below.

### 4.2 Behavioural experiments

#### 4.2.1 Animals

Third-instar feeding-stage larvae (*Drosophila melanogaster*), aged 5 days after egg laying, were used throughout. Flies were maintained on standard medium, in mass culture at 25 °C, 60-70 % relative humidity and a 12/12 hour light/dark cycle. We took a spoonful of food medium from a food vial, randomly selected the desired number of larvae, briefly rinsed them in tap water, and started the experiment.

For the experiments regarding the Cirl mutant, we used two genotypes: the *Cirl*^*KO*^ null mutant and a genomic rescue strain, *Cirl*^*Rescue*^. For a detailed description of the genotypes, see [40].

For optogenetic experiments, we used transgenic larvae to express the ChR2-XXL light-gated ion channel in DANs. To this end, the effector strain *UAS-ChR2-XXL* (Bloomington Stock Center no. 58374) [43] was crossed to one of two driver strains: either *TH-Gal4* (Bloomington Stock Center no. 8848) or *R53F07-Gal4* (Bloomington Stock Center no. 50442) to obtain double-heterozygous offspring. As driver controls, the driver strains were crossed to a local copy of *w*^*1118*^ (Bloomington Stock Center no. 3605, 5905, 6326). As effector controls, either *w*^*1118*^, or a strain carrying the landing site used for the Gal4 (attP2), yet without a Gal4 domain inserted (“empty”) [80], was crossed to *UAS-ChR2-XXL*. Because ChR2-XXL is sensitive enough to be activated by daylight (not shown), the flies were raised in vials constantly darkened by black cardboard wrapping.

#### 4.2.2 *Cirl*^*KO*^ experiments

For the experiments regarding the Cirl mutant, the larvae were placed in the centre of Petri dishes of 9 cm inner diameter (Sarstedt, Nümbrecht, Germany), filled with 1 % agarose (Biozym Scientific GmbH, Hessisch Oldendorf, Germany). The Petri dish was placed in a custom-built arena lit up with infrared light (PeakTech DC Power Supply 6080). Always five larvae were placed on a Petri dish, and their behaviour was recorded for 1 minute with a camera (Logitech C920 HD Pro).

#### 4.2.3 Optogenetic activation experiments

For optogenetic experiments, the larvae were placed on Petri dishes of 9 cm inner diameter, filled with 1 % agarose. Experiments were performed inside a 43 × 43 × 73 cm surrounding box equipped with a custom-made light table featuring a 24 × 12 LED array (470 nm; Solarox, Dessau-Roßlau, Germany) and a 6 mm thick diffusion plate of frosted plexiglass on top to ensure uniform blue light for ChR2-XXL activation (120 µW/cm^2^). The Petri dishes were placed directly on top of the diffusion plate. They were surrounded by a polyethylene diffusion ring; behind the diffusion ring 30 infrared LEDs (850 nm; Solarox, Dessau-Roßlau, Germany) were mounted to provide illumination that was invisible to the larvae, yet allowed the recording and tracking of their behaviour for offline analysis. To this end, a camera (Basler acA204090umNIR; Basler, Ahrensburg, Germany) equipped with an infrared-pass filter was placed above the Petri dish.

Experiments were performed by placing seven to eight larvae in the middle of a Petri dish, and letting them walk freely for two or four minutes. The first 30 s were in darkness, followed either by a 30 s light stimulation period and another 60 s of darkness, or by three cycles of 10 s light stimulation and 60 s of darkness.

### 4.3 Tracking

We developed the IMBAtracker to track multiple larvae on a single Petri dish while keeping track of their identity. Although several tracking solutions have been developed in recent years [17–27, 31], the development of new tracking software seemed necessary to ensure two important aspects: first, increasing and controlling the overall measurement precision according to the needs of the analysis; second, reducing the data fragmentation caused by collisions or by the larvae leaving the region of interest. This attempt to reduce fragmentation is especially important since it paves the way for the individual-level analysis.

#### 4.3.1 Architecture

The tracker is designed to process a region of interest (the Petri dish area), which is defined per video. We do not require the region to contain all the objects of interest at all times (i.e. larvae can exit the region of interest). Videos should be made from above or below with a minimum resolution of 30 pixels per animal (although higher resolutions improve accuracy). Either a light background with dark objects or a dark background with light objects is supported, and any colour information is discarded as the image is converted to greyscale. In order to deal efficiently with the issue of fragmentation, it is crucial to separate the tracking into three distinct phases (see below). The general architecture is depicted in Fig. 13.

#### 4.3.2 Implementation

The software is written in C++ using OpenCV [81] as a base for input-output and image processing, lpsolve for linear optimization, and further standard algorithms and data structures. This ensures fast processing of the videos, which can be performed on a commodity PC. A simple user interface (written in Python) is provided.

#### 4.3.3 First Phase

The first tracking phase works similarly to other tracking software: we perform image processing, segmentation and extraction of objects of interest as connected components. However, instead of discarding objects during collisions, the contours are used to detect and record the types and objects involved in those collisions. An overview of the process can be seen in Fig. 13A.

We perform the following image-processing steps to extract the foreground from the background and to separate our objects of interest:

1. **Background subtraction**: We opt for a very simple background subtraction. The background model consists simply of the pixel-wise average value of 100 equidistantly spaced frames of the central third of the video to increase robustness against sudden video changes towards the beginning or end of the video.
2. **Thresholding**: A threshold separates the image into two classes: the objects of interest in the fore-ground, and the background. To perform this thresholding in an automated fashion we make use of Otsu’s method [82], which selects the threshold that minimizes the in-class variance.
3. **Blob extraction**: This step tries to extract all connected components of the foreground as objects, which we refer to as blobs. For this step we use a library called cvBlob. The algorithm used by the library is derived from [83].

At each video frame we receive new blobs from the blob extraction routine. In order to see how well the blobs match the blobs from the previous frame, we test each blob from the new frame on its overlap with all blobs from the previous frame. Here we have the following categories:

- 1 − 1: 1 blob from the old frame matches 1 blob from the new frame.
- 1 ∼ 1: 1 blob from the old frame partially matches 1 blob from the new frame.
- 1 − *N* : 1 blob from the old frame matches *N* blobs from the new frame.
- *N* − 1: *N* blobs from the old frame match 1 blob from the new frame.
- *N* − *M* : *N* blobs from the old frame match *M* blobs from the new frame.

The latter three cases point to collisions of larvae, as they occur, for example, when larvae merge with one object from a given frame (N-1), as well as when they split again (1-N). In this way, blobs that consist of several larvae can be separated from blobs that probably only consist of a single larva. This information can be visualized in a collision graph with parental nodes (blobs that split into several new blobs) and child nodes (blobs that emerge from the parental node) that will be used in the second phase of the tracking.

For those blobs that could be single larvae we try to extract some key information on the shape. To do this, we first reconstruct the contour using parametric cubic smoothing splines. We generate a contour of 200 points for each blob based on the splines. We then compute curvatures of the reconstructed contour points and, after applying a multiple-pass moving average filter over the contour for noise reduction, we detect the two biggest local maxima. These indicate the endpoints (head, tail) of the larva and are the base for the spine reconstruction. We always match the endpoints across frames based on distance, thereby storing the trajectories of each endpoint during the track. For cases where the continuation is unclear (such as where there is curling or a spine cannot be reconstructed), the track is interrupted and split into subtracks in which the endpoints’ trajectories are continuous.

#### 4.3.4 Second Phase

After the basic processing of the video, we attempt to clarify and reduce the fragmentation of the data by merging subtracks. We initially work on two aspects, the larva count in each blob and accurate head/tail detection. We then tackle the issue of fragmentation.

To clarify whether a blob is a larva or a cluster of colliding larvae, we observe that a good approximation of the contents is given by trying to find the minimum number of larvae needed to fulfil the parent-child constraints, namely that parental nodes have the same number of larvae as their children nodes. We formalize this as a linear optimization problem over a collision graph. Nodes store the uncertain number of colliding larvae, which is to be determined during optimization, and edges represent the parent-child relationship.

Given the collision graph *G*(*V, E*), where *V* are the nodes and *E* the set of edges (*i, j*) from node *i* to node *j*, we want to solve the following integer optimization problem

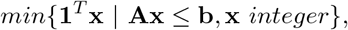

where vector **x** = (*x*_1_, …, *x*_*n*_) contains the number of larvae for every node *i* as components *x*_*i*_, vector **1** = (1, …, 1) *∈* {1}^*n*^, and the system of inequalities **Ax** *≤* **b** is constructed from the following two constraints with the sets of parent nodes *P*_*i*_ and child nodes *C*_*i*_ of a node *i*.

The number of larvae *x*_*i*_ in a blob *i* is at least the number of incoming parent edges, outgoing child edges or 1, whichever is highest:

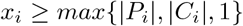

The number of parent larvae and the number of child larvae of every blob, more specifically every internal node *i*, should be in balance:

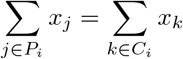

Note that this equality constraint can be rewritten as two non-strict inequalities for the system of inequalities in the above minimization problem.

Matrix **A** is totally unimodular, which guarantees that the problem is tractable, that we can use linear optimization techniques to solve it, and that these techniques will return integral solutions only [84].

We can now proceed to disambiguate the head/tail assignment. As mentioned above, all larva tracks are split into sub-tracks in which the endpoints have clear and continuous trajectories. For these trajectories we calculate several metrics per endpoint (Fig. 13B, middle panel), such as:

- Average brightness around 10 pixels of the endpoint
- Total distance covered
- Total distance of the endpoint relative to the centroid in the previous frame, i.e. the endpoint moves towards or away from the centroid.

The head is then assigned according to votes from each of these measurements. We assign one head vote to endpoint A and one tail vote to endpoint B in each of the following cases (and the reverse otherwise):

- Average brightness of A *<* Average brightness of B
- Total distance covered of A *>* Total distance covered of B
- A moves away from the centroid

We assign endpoint A to be the head if it has more head votes than endpoint B; otherwise we assign B to be the head.

As our main interest lies in studying the behaviour of freely moving larvae, we do not try to disentangle their behaviour within a collision. In order to decrease the fragmentation of tracks, we decided first to disentangle the ephemeral collisions. During these, behaviour should be minimally affected, and their elimination should allow for longer tracks and greater preservation of identity.

We define a larva shape model for tracking purposes based on orientation and four joint angles as depicted in Fig. 13B (right-most panel). In order to avoid wrong assignments, we restrict this larva model to ephemeral collisions, i.e. collisions of two larvae that last less than eight seconds, or short touches of single larvae on larger blobs. This representation is initialized for the collision participants one frame before the collision, using the values that we have already calculated. For each frame of the collision, the contour of each larva in collision is approximated by subtracting the contours of the counterparts from the previous frame from the complete contour of the current frame. We then search, using a brute-force grid search, through the parameter space (i.e. orientation and joint angles) for the best neighbouring fit of our model to the approximated contour. When the collision is finished we compare the centroids of the collided models of the last frame of the collision to the new larvae coming out of it and assign the closest match.

#### 4.3.5 Third Phase: Statistical Collision Resolution

The first approach for resolving collisions, described above, is limited to collisions of two larvae that do not take longer than 8 s. For collisions that involve up to three larvae, or take longer, a second approach for collision resolution based on statistical attributes is adopted. In brief, for each larva entering a collision that was not already resolved by the first approach, the distributions of its body length, width, area and perimeter values are compared by non-parametric testing with those of each larva leaving the collision (Fig. 13C).

Taking a collision of two larvae as an example, we first perform a principal component analysis (PCA) over the standardized values of the body length, width, area and perimeter, using only the first resulting principal component, which explains about 83 to 94 % of all variance in a test data set, for all further steps. Then we perform Mann-Whitney U-tests between the two ingoing and the two outgoing larvae, and compare the resulting p-values. The rationale is that a high p-value indicates little difference between the data, so the higher the p-value, the higher the chance that an ingoing and an outgoing larva are actually the same individual. Such comparison is legitimate if all the tests include the identical sample size [85]. Therefore, we restrict the tests to the minimal number of frames for any larva involved (i.e. the frames before the collision for the ingoing larvae, and the frames after the collision for the outgoing larvae), up to a maximum of 45 frames, as higher frame numbers decreased the rate of correct assignments in a test data set (not shown). With two ingoing (larvae 1 and 2) and two outgoing (larvae A and B) animals, there are only two logical assignments: either 1-A and 2-B is correct, or 1-B and 2-A. A collision is resolved if both p-values for one assignment are higher by a factor of 1000 (i.e. the larvae are more similar) than both p-values for the other assignment. If no logical assignment can be made, or the difference between the p-values is too small, the collision remains unresolved. Collisions of three larvae are performed in an analogous way. We do not attempt to resolve collisions of four or more animals, as the chances of a resolution are extremely low.

We also apply this approach to asymmetrical collisions, e.g. when a single larva leaves a collision of three animals or when the tracker loses track of larvae either at the Petri dish wall or at objects such as odour containers in olfactory experiments. In the latter cases, whenever a larva leaves the Petri dish wall, for example, it is statistically compared to all larvae that were previously lost at the wall. In these asymmetrical cases, we perform additional U-tests between all ingoing larvae (e.g. all larvae that have been lost at the wall). Such a collision is resolved if the p-value of a test between the outgoing and one of the ingoing larvae is higher by a factor of 1000 than all other p-values. If no logical assignment can be made, or the difference between the p-values is too small, the collision remains unresolved.

### 4.4 Data Analysis

The main output of the IMBAtracker is a time series of 12 spine points that reach from the head of the animal to its tail, as well as 24 contour points (Fig. 1). These data are the basis for all further data analyses and visualisations with the IMBAvisualiser. To prevent multiple tracks from the same individual animal from entering the analysis, we discarded all tracks shorter than half of the video length after the collision resolution.

The spatial resolution in our video recordings was not sufficient to resolve the peristaltic wave along individual body segments. However, we noticed that the rhythmic velocity changes of the tail point provided a reliable indicator for the occurrence of one peristaltic forward movement (Fig. 1). We therefore defined each local maximum in the tail forward velocity as one step, and the period between two maxima, i.e. one complete peristaltic cycle, as an inter-step (IS) period (Fig. 1D). Since the local-maxima detection algorithm we employed also produced false maxima during intervals of little or no forward motion, we discarded maxima detected whenever the tail speed in a forward direction was less than 0.6 mm/s.

Lateral head casts are a very prominent feature during larval chemotaxis on a plain agarose surface. We quantified these head movements based on the angular speed during the head vector, where the head vector was defined as the vector between the ninth and the eleventh spine point (Fig. 1B). In order to detect head casts to the left (or right) we first detected all intervals of angular speed larger than 35°/s (or smaller than −35°/s) (Fig. 1C). The start and end points of these intervals correspond to salient movements of the head part of the animal to the left or right side. We next introduced two criteria in order to discard incorrectly assigned head casts. (1) We discarded any initially detected head cast whenever the angular speed of the tail vector during head casting exceeded 45°/s. This criterion takes account of the visual observation that whenever the tail parts of the animal reorient, these periods most likely do not coincide with an active lateral head casting movement, but rather a stretching out of the body bending angle when stepping is resumed. (2) We discarded any initially detected head cast whenever more than one step was detected during head casting. This criterion takes account of the observation that any active lateral head casting interval can coincide with at most one forward stepping detection point.

After the detection of runs, steps and head casts, we calculated a number of behavioural attributes. In the following we provide a detailed description of all the attributes used in this study. A full list of attributes implemented by the IMBAvisualiser is available online (https://github.com/XXX). Please note that for some attributes two versions exist, using either mm or individual body lengths (bl) as the unit. Unless explicitly mentioned in the results, it was always the version using body lengths that was applied.

#### 4.4.1 Attributes

Whenever the behaviour of an individual larva is displayed over time (Fig. 1, 10-E1), the respective attributes are calculated per video frame. Whenever the average behaviour of a group of larvae is displayed over time (Fig. 7B, 7-E1, 8A,C,E,G, 9B, 9-E1, 11, 12), the mean and the 95 % confidence intervals of the respective attribute are calculated per 2 s time bin. Whenever data are displayed in box plots to compare across experimental conditions, or in scatter plots to determine correlations, the mean of each attribute is calculated for each individual larva across the whole observation period.

For the optogenetic experiments, we were interested in the difference in a given attribute X between different time intervals. In these cases we calculated Δ-values as the difference in X during each of the time intervals. For Figures 7 and 9, we calculated X during the 30 s of light stimulation minus X during the 30 s before light stimulation. For Figure 8, we calculated X during the 10 s of light stimulation minus X during the 10 s before. In Figure 12, we calculated X in the 10 s after a switching point minus X in the 10 s before the switching point. In Figure 12, we additionally calculated so-called peak values of the absolute bending angle and the absolute HV angular speed in order to quantify the local maxima of these values at the forward switching points. The peak values were calculated as X during the 10 s interval around the switching point minus X during the 5 s before and the 5 s after that interval.

#### 4.4.2 User Interface for Visual Analysis

The interactive IBMAvisualizer is a visual analytics concept [86]. Visual analytics generally enculapses approaches such as visually supported dimension reduction techniques [87–89] or visual paramaterization support [90, 91] in order to support a human-centered context-related analysis cycle [92]. The IMBAvisualiser was created using the R programming language [93] and the Shiny library, which allows user interfaces to be built for data visualization [94]. For this, the analysed data sets are saved in RDS files which are serialized R objects. These RDS files can be uploaded in order to plot various visualizations based on the attributes that are calculated. In addition, the data are saved in CSV files that can easily be opened and further processed by other software. In the following paragraphs, we very briefly introduce the essential functions of the software. A detailed documentation can be found online along with the code.

In the “Single Mode” the user has the possibility to visualize individual tracks, compare different time series attributes, and compare different tracks. The track selection menu allows multiple tracks to be selected. This way of analysing the data permits a detailed analysis of the time dependent attributes for individual animals. This detailed analysis of individual animals’ time series allowed us to find anomalies in the behavior of the larvae.

Data aggregation allows attributes to be calculated, usually as means or variance, from all the data either on the Petri dish or the individual animal. Using different sliders, it is possible to filter the aggregated data by time, absolute bearing angle, absolute HC angle, and the distance from the source of a stimulus, e.g. an odour (in experiments using such a stimulus). For the analysis of backwards-walking larvae, it is also possible to filter the data based on the run direction (forwards, backwards). These filtering options permit a detailed analysis for different subsets of the data.

We used random forest models, which were established by Leo Breiman and are based on CART (Classification and Regression Trees) [95, 96]. The random forest algorithm works by building N number of decision trees and then predicting the class label of an instance using a majority voting. Each tree is trained on a random subspace of the data, only using a sample of the instances. To evaluate the random forest classification, we used five-fold cross validation, which was repeated three times. Five-fold cross validation means that the test set for the model is used from five different, disjunct sets of the data set. The data set is then shuffled, and the procedure is repeated three times. Cross validation is used to make sure that the model is not based merely on the selected data, but can explain the full data set.

For the applications of the random forest in this study, we pre-selected 30 behavioural attributes to be included (Tables 2-3). The pre-selection was warranted to avoid using largely redundant attributes (such as the IS speed in both *bl/s* and in *mm/s*), and to exclude non-behavioural attributes such as the size of the larvae. In order to provide a good estimate of the importance of each attribute, the complete random forest model (including the cross validation) was performed 10 times for any given data set, and the importance rank of each attribute was noted.

### 4.5 Statistical Tests

Two-tailed non-parametric tests were used throughout. When multiple comparisons were performed within one analysis, a Bonferroni-Holm correction was applied to keep the experiment-wide error rate below 5 % [97]. Values were compared across multiple independent groups with Kruskal-Wallis tests (KW tests). In case of significance, subsequent pair-wise comparisons used Mann-Whitney U-tests (MW tests). In cases of within-animal comparisons, as well as for comparison to a single baseline (Fig. 2G) we used Wilcoxon signed-rank tests (WS tests). For correlations, we employed Spearman’s rank correlation test (SC tests).

## Supporting information

Rich media file 1

Rich media file 2

Rich media file 3

Rich media file 4

## 5 Acknowledgements

This study received institutional support from the Leibniz Institute for Neurobiology, the Wissenschaftsgemeinschaft Gottfried Wilhelm Leibniz, and the University of Leipzig; as well as grant support from the Deutsche Forschungsgemeinschaft (DFG) (to B.G. and M.S.: GE 1091/4-1; to B.G.: FOR 2705; to N.S.: FOR 2149, SCHO1791/1-2, CRC 1423), the European Commission grant MINIMAL (to B.G.: FP7 – 618045) and the German-Israeli Foundation for Science (GIF) (to M.S.: G-2502-418.13/2018). Technical assistance by M. Dombach, F. Unterstab and the Combinatorial NeuroImaging Core Facility at the Leibniz Institute for Neurobiology, as well as discussions with C. König, M. Nawrot, P. Sakagiannis, J. Thoener, N. Toshima and B. Webb are gratefully acknowledged. We thank R.D.V. Glasgow (Zaragoza, Spain) for language editing.

## 7 Author contributions

Conceptualization: M.S., M.T., E.P., B.G. Methodology: M.T., E.P., T.S., S.G., D.J.L., M.S. Software: M.T., E.P., T.S., S.G., A.-R.K. Formal analysis: M.T., M.S., A.-R.K. Investigation: M.T., A.-K.D., A.-R.K., M.S. Data curation: M.T., A.-K.D., A.-R.K., M.S. Writing - original draft: M.S., M.T., T.S., E.P. Writing - review & editing: M.T., E.P., T.S., A.-R.K., S.G., A.-K.D., N.S., B.G., D.J.L., M.S. Visualization: M.T., M.S., E.P., T.S., D.J.L. Supervision: M.S., B.G., N.S., D.J.L. Project administration: M.S. Funding acquisition: B.G., N.S., D.J.L., M.S.

## 8 Extended Figures

**Fig. 1-E1:**
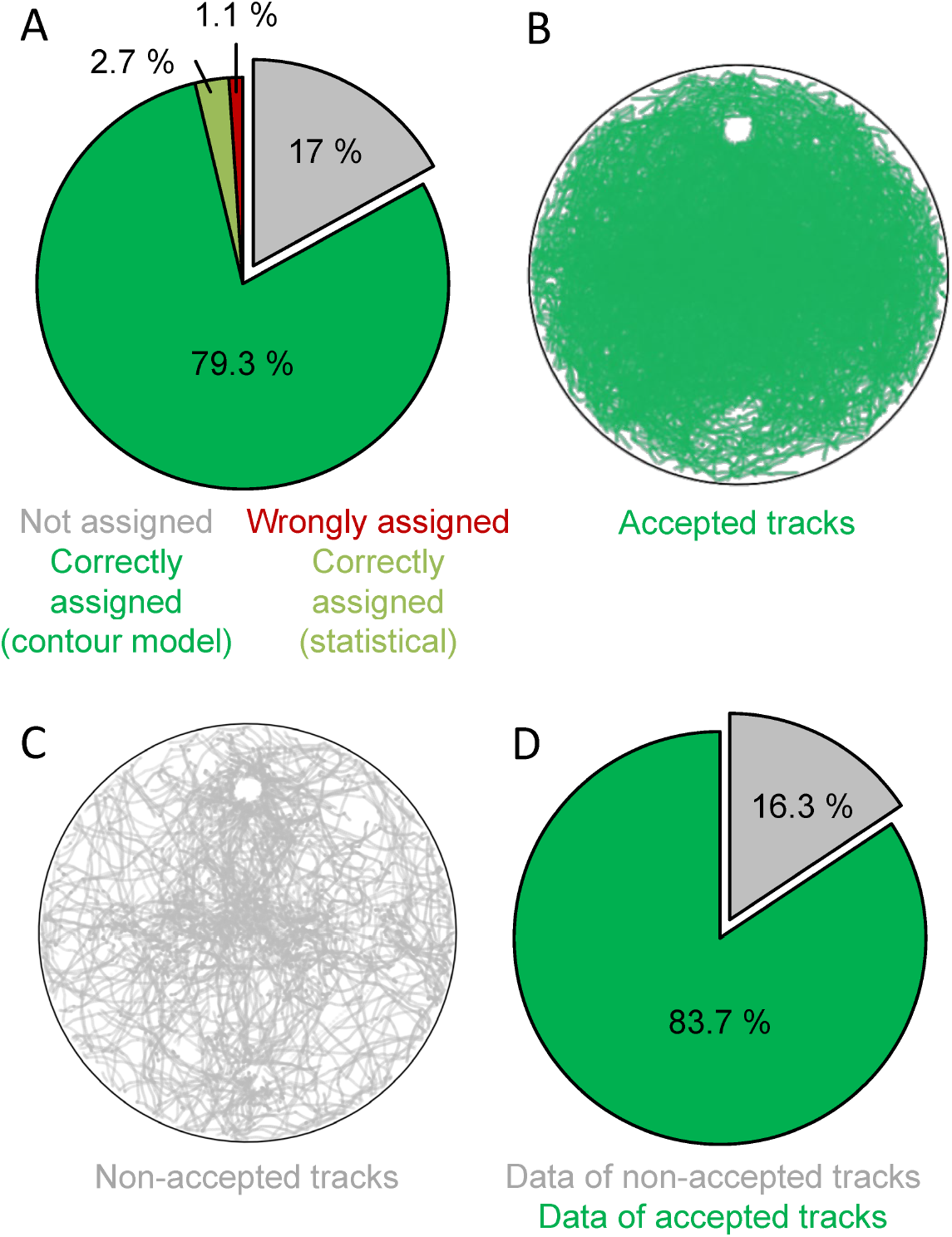
Evaluation of the collision resolution. (A) Manual evaluation of all collisions between two larvae in the data set. We attempted to resolve 83 % of all collisions, and 99 % of those were correct. (B) All tracks of individuals accepted for the analysis. We accepted only larvae that were observed for at least half of the duration of the recording (in this case, 90 s). Two areas are free from tracks as they were covered by teflon containers for odour application. (C) All tracks not considered for the analysis. (D) Percentage of all data points of the accepted and non-accepted tracks.

**Fig. 1-E2:**
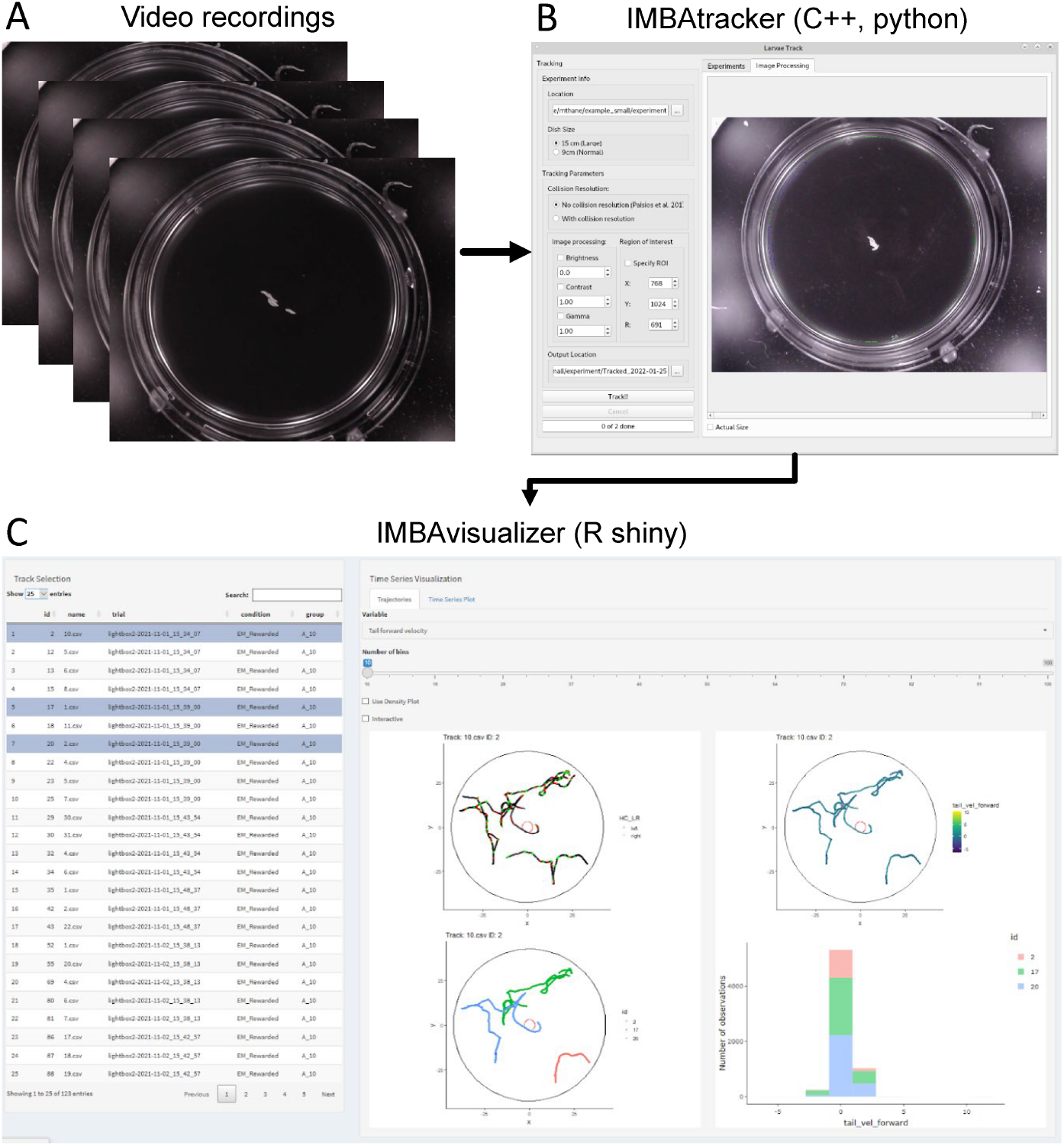
The individual maggot behaviour analyser (IMBA). (A) Video recordings can be obtained from various sources, either providing bright objects on a black background, or dark objects on a white background. (B) The IMBAtracker (written in C++, controlled by a graphical user interface, written in Python) identifies larvae with their spines, contours, heads and tails, resolves collisions, and provides coordinates for each spine and contour point in each frame for every larva detected. (C) The IMBAvisualiser (written in R shiny) allows flexible visualization and analysis of the tracked data. This includes calculating 95 behavioural attributes, performing machine-learning algorithms and basic statistical tests, displaying the behaviour of individual larvae, and quantifying large data sets with a multitude of options.

**Fig. 2-E1:**
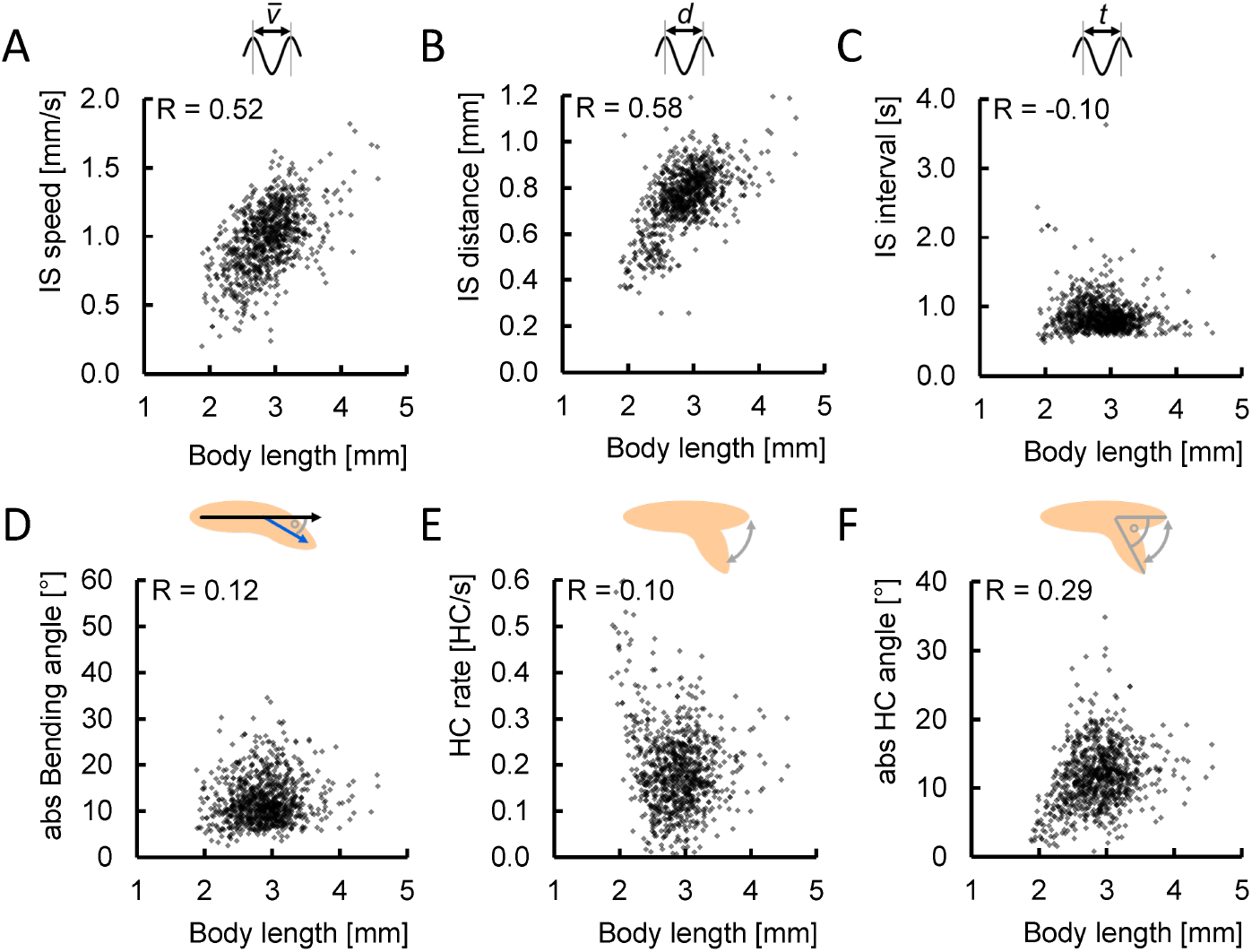
Crawling speed of larvae is correlated with body length. Of the six basic locomotor attributes, we found (A) the IS speed and (B) the IS distance to be mildly correlated with the individual animals’ body length. In contrast, no correlations were found between body length and (C) the IS interval, (D) the absolute bending angle, (E) the HC rate, or (F) the absolute HC angle. Correlations are determined by SC tests. The underlying source data, as well as the results of the statistical tests, can be accessed in the “Figure 2-source data” file.

**Fig. 4-E1:**
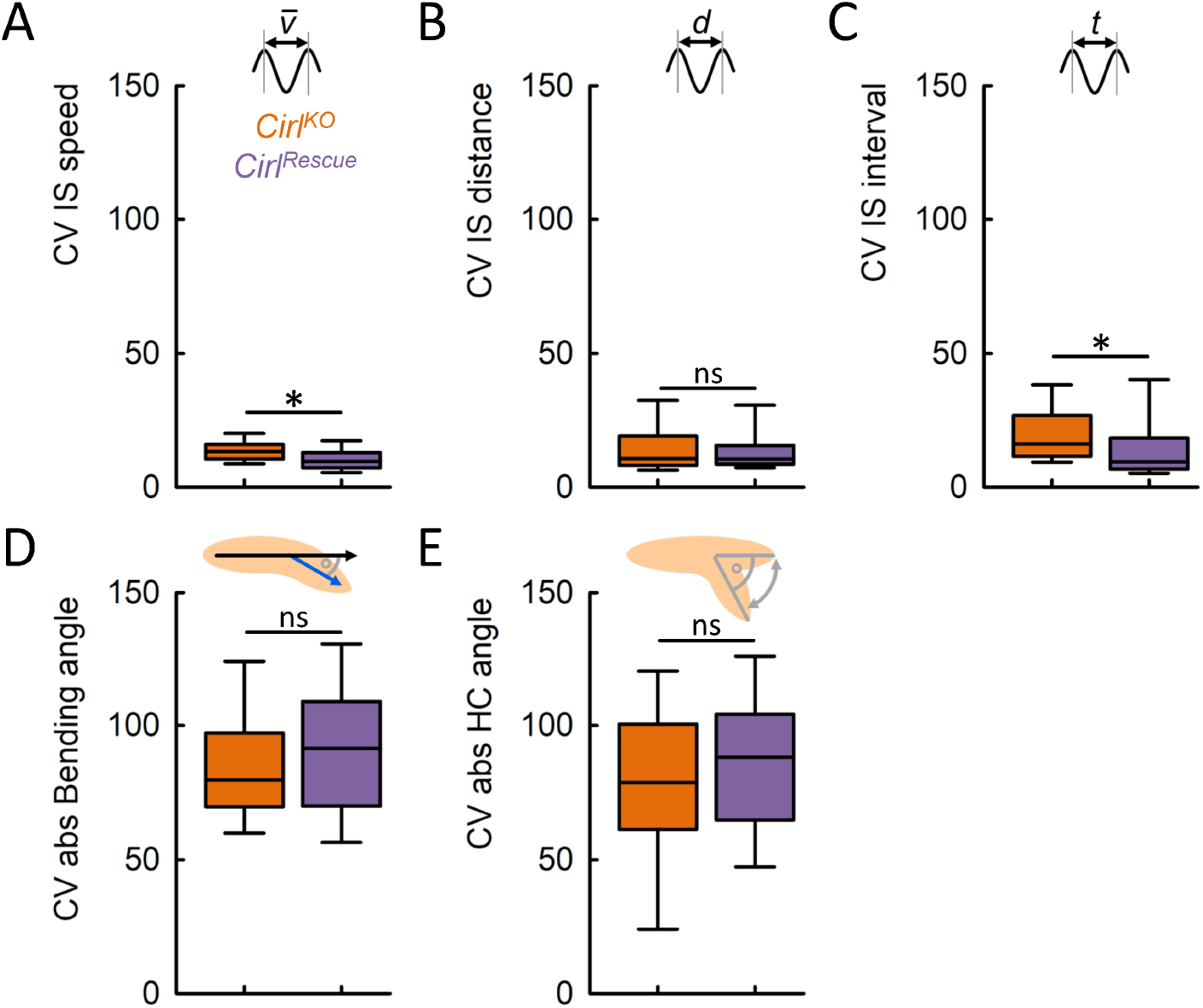
Intra-individual variability of speed and duration of peristaltic forward cycles is increased in the *Cirl*^*KO*^ mutant. We determined the coefficient of variation (CV) for each basic locomotor attribute (except the HC rate) in each individual animal, and found (A) an increased variability for the IS speed, (B) no change for the IS distance, (C) increased variability for the IS interval, and no change for (D) the absolute Bending angle and the absolute HC angle. Displayed are the median as the middle line, the 25 % and 75 % quantiles as boxes, and the 10 % and 90 % quantiles as whiskers. Asterisks indicate significant differences in MW tests. The underlying source data, as well as the results of the statistical tests, can be accessed in the “Figure 4-source data” file.

**Fig. 7-E1:**
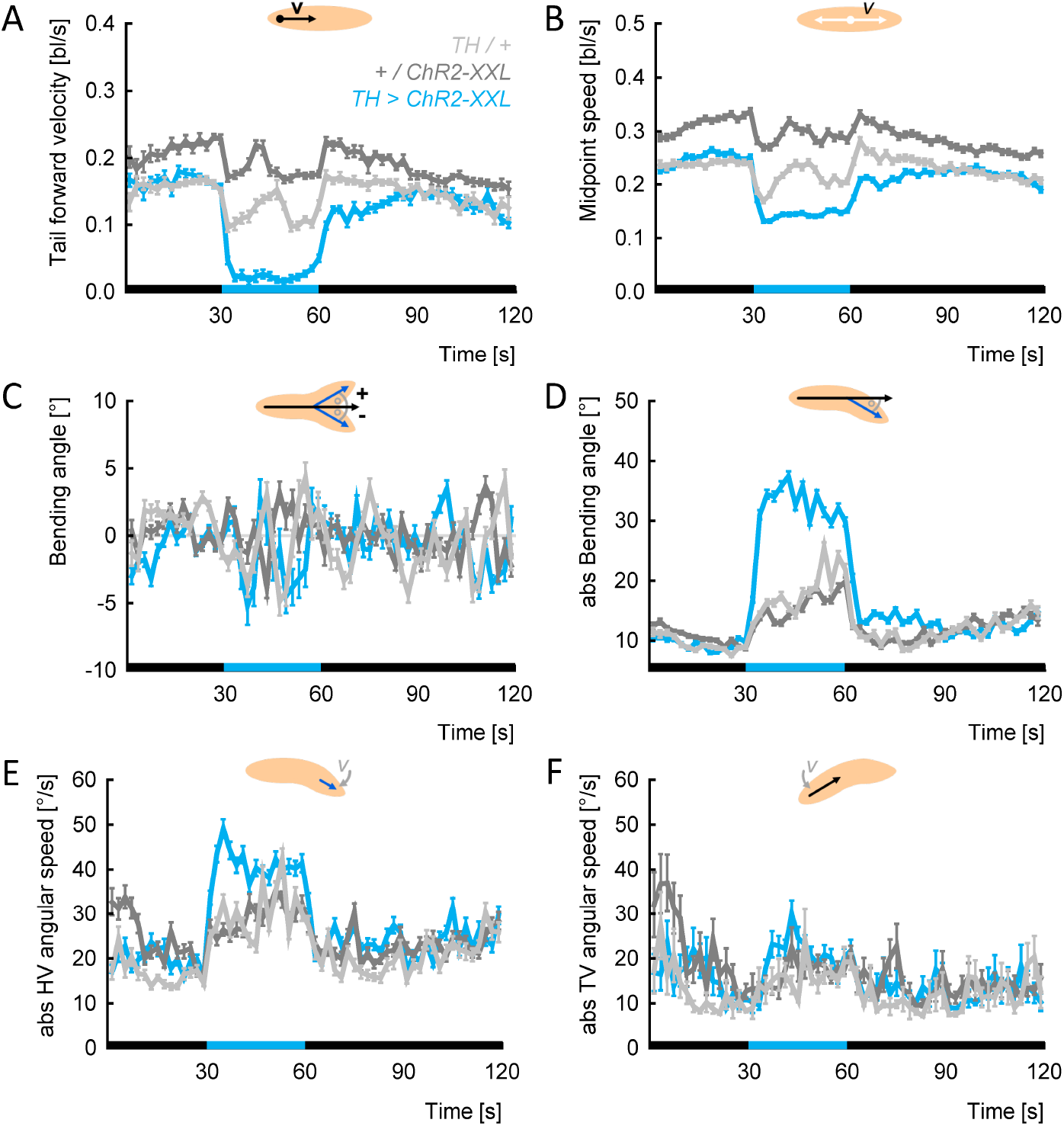
Dopamine neuron activation reduces speed and increases bending. Displayed are the mean and 95 % confidence intervals of all the data for each genotype within each 2 s bin. Black and blue stripes on the X-axis indicate periods of darkness and blue light stimulation, respectively. Of the ten behavioural attributes found in Fig. 7, the travelled distance was omitted as it is a cumulative measure, and the measures of intra-individual variability were omitted as this kind of display does not take into account individual behaviour. (A) Tail forward velocity, (B) midpoint speed, (C) bending angle, (D) absolute bending angle, (E) absolute HV angular speed, (F) absolute TV angular speed. The underlying source data can be accessed in the “Figure 7-source data” file.

**Fig. 8-E1:**
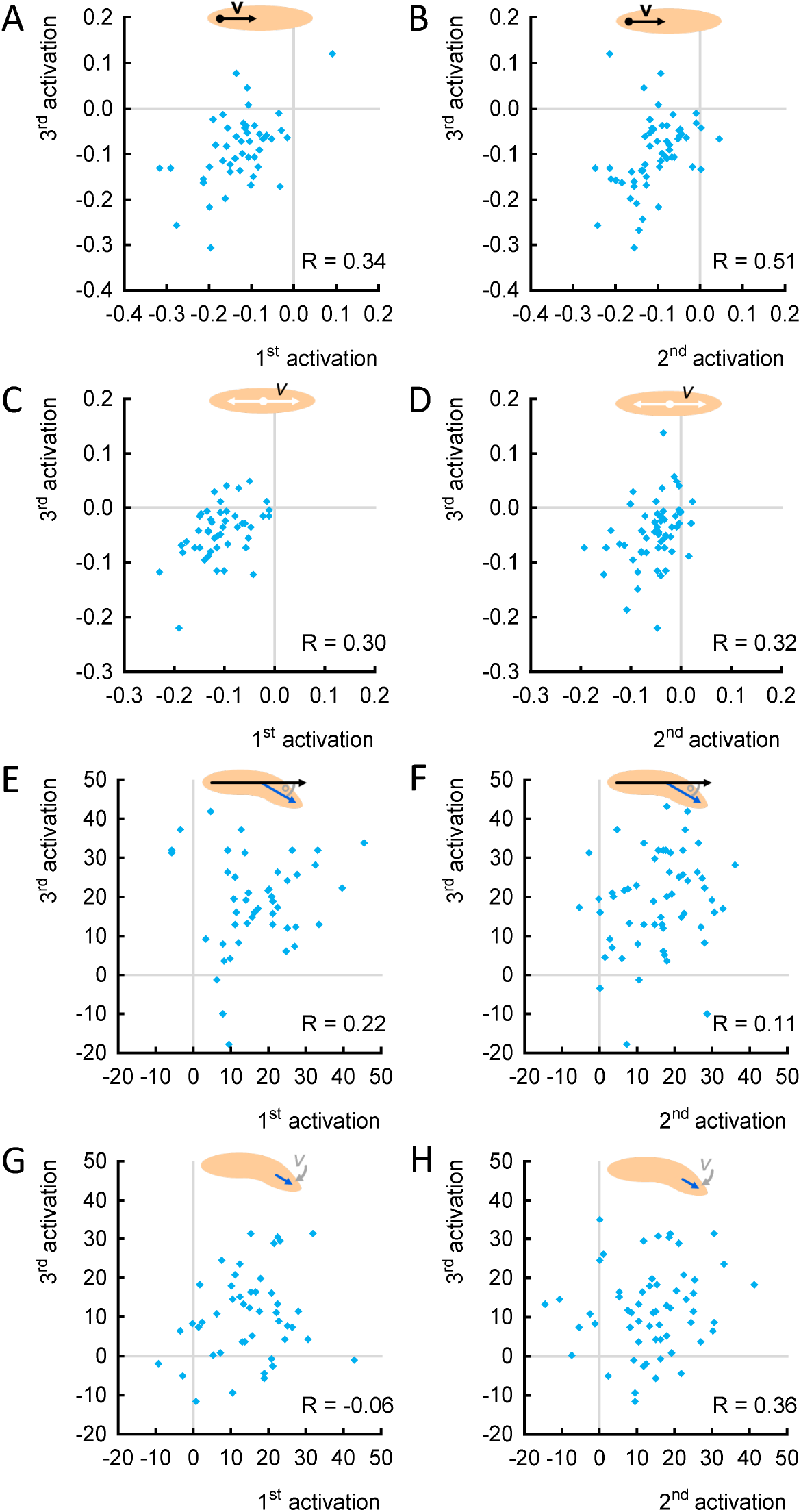
Individuals reduce speed consistently upon repeated optogenetic activation of dopamine neurons. (A-B) The tail forward velocity during light stimulation was normalized to each individual’s behaviour in the 10 s before light activation (called Δ-values). The Δ-values of (A) the first and third activation, as well as (B) the second and third activation, were positively correlated. (C-D) As in (A-B), but for the midpoint speed. (E-F) As in (A-B), but for the absolute bending angle. (G-H) As in (A-B), but for the HV absolute angular speed. Correlations were determined by SC tests. The underlying source data, as well as the results of the statistical tests, can be accessed in the “Figure 8-source data” file.

**Fig. 9-E1:**
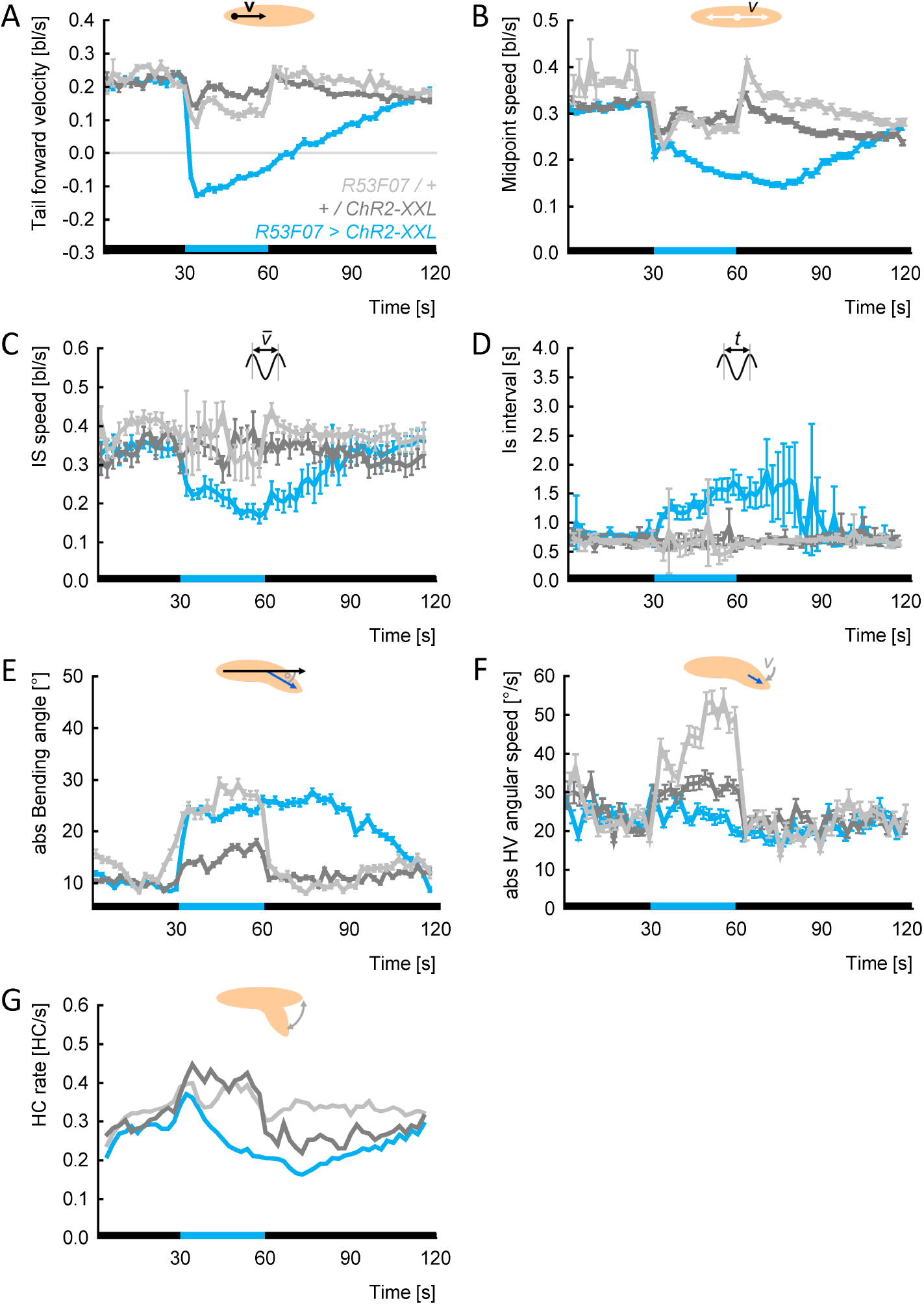
Activation of *R53F07-Gal4* neurons triggers backward locomotion, reduces speed and increases bending. Displayed are the mean and 95 % cconfidence intervals of all data for each genotype within each 2 s bin, except in case of the HC rate, which was calculated within each 2 s bin by dividing all frames with a HC by all the frames in the bin. Black and blue stripes on the X-axis indicate periods of darkness and blue light stimulation, respectively. Of the ten behavioural attributes found in Fig. 8, the travelled distance was omitted as it is a cumulative measure, and the measures of intra-individual variability were omitted as this kind of display does not take into account individual behaviour. (A) Tail forward velocity, (B) midpoint speed, (C) IS speed, (D) absolute bending angle, (E) HV absolute angular speed, (F) HC rate. The underlying source data can be accessed in the “Figure 9-source data” file.

**Fig. 10-E1:**
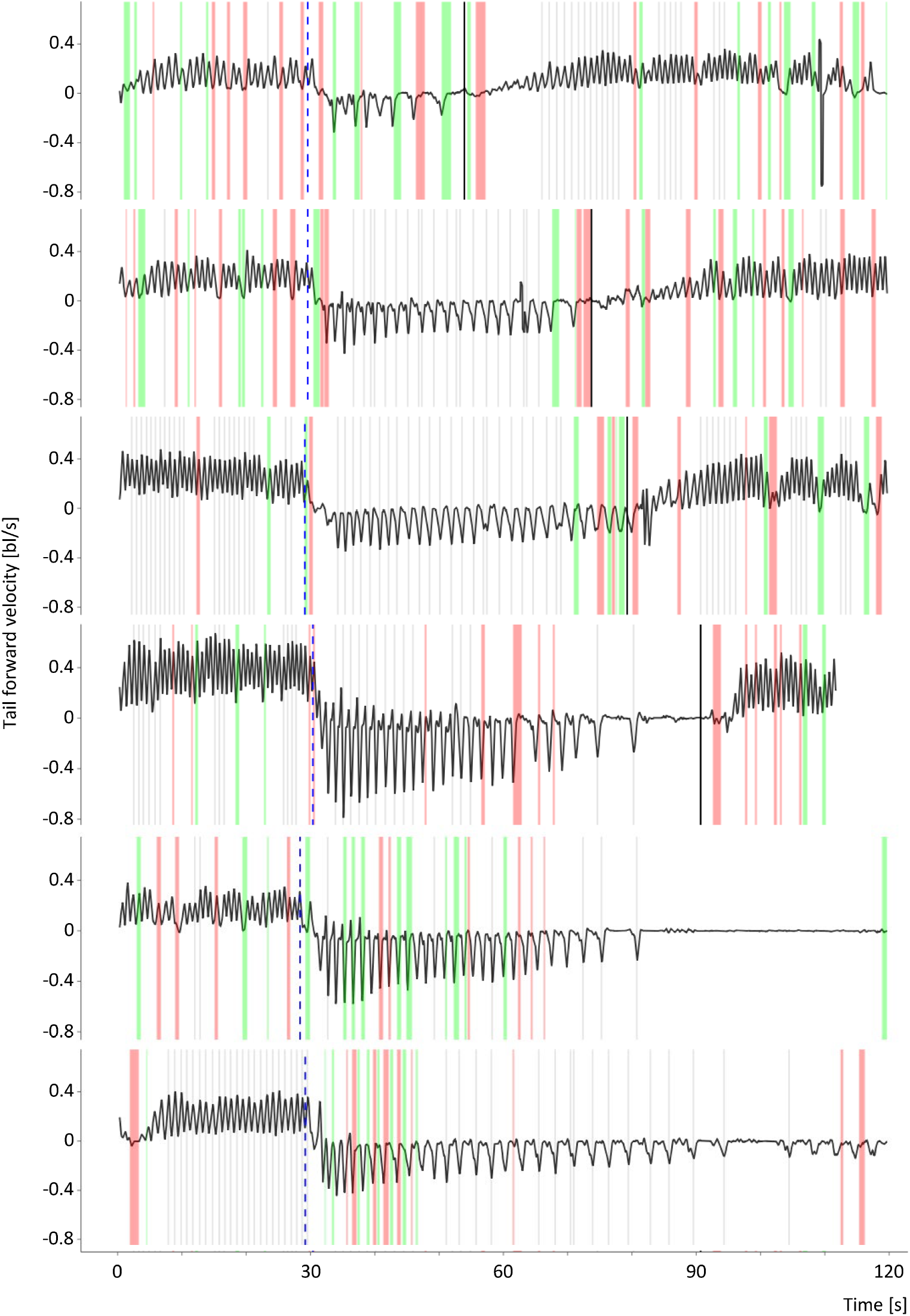
Velocity of sample individuals over time. Six sample animals of the experimental genotype (*R53F07 >ChR2-XXL*) were picked to display the tail forward velocity over the course of the experiment. Blue light stimulation started at 30 s and ended at 60 s. Stippled blue lines indicate the automatically detected switch from forward to backward locomotion, and solid black lines the switch from backward to forward locomotion. The two animals at the bottom had not reverted to forward locomotion by the end of the recording. For other details, see Fig. 1.

**Fig. 12-E1:**
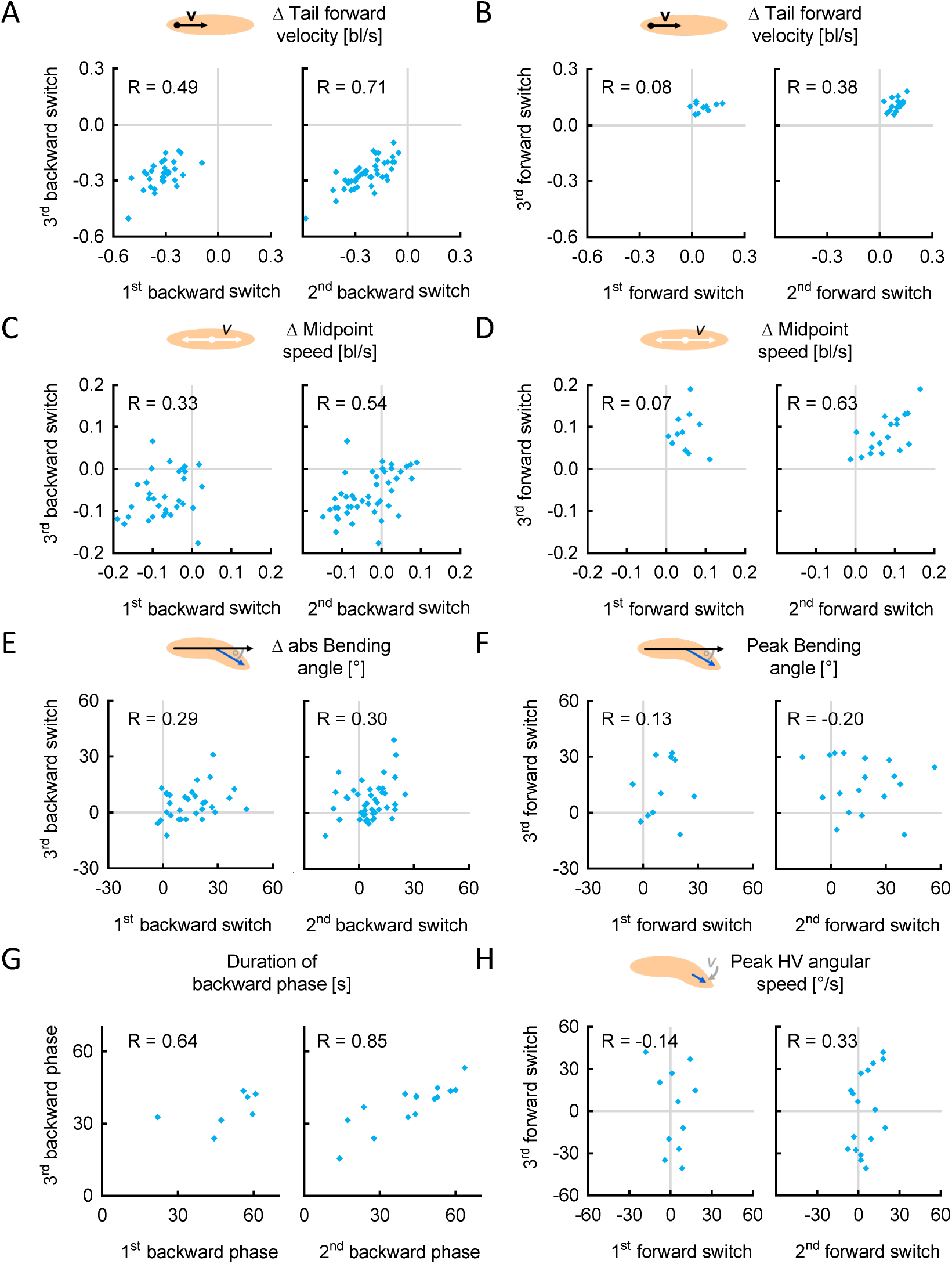
Individuals change speed consistently upon repeated backward, but not forward switches. Individual data were aligned either to the switch from forward to backward crawling (A,C,E), or to the switch from backward to forward crawling (B,D,F,H). (A) The tail forward velocity during the 10 s after the switch was normalized to each individual’s behaviour in the 10 s before the switch. These Δ-velocities of (left) the first and third activation, as well as (right) the second and third activation, were positively correlated. (B) As in (A), but for the forward switches. (C-D) As in (A-B), but for the midpoint speed. (E) As in (A), but for the absolute bending angle. (F) To quantify the peak of the absolute bending angle at forward switches, we normalized the behaviour in the 10 s around the switch with the 5 s before and the 5 s after (called peak values). (G) The durations of the first and third, as well as the second and third, backward crawling phases were positively correlated.(H) As in (F), but for the HV absolute angular speed. Correlations were determined by SC tests. The underlying source data, as well as the results of the statistical tests, can be accessed in the “Figure 12-source data” file.

## 9 Rich media files and source data files

### 9.1 Rich media files

**Rich media file 1**: Animation of a sample *Cirl*^*Rescue*^ control larva. The blue dot marks the position of the head. The filling indicates the tail forward velocity in [bl/s]. The fast and regular peristaltic cycles can be easily seen by the quick colour changes.

**Rich media file 2**: Animation of a sample *Cirl*^*KO*^ mutant larva. The blue dot marks the position of the head. The filling indicates the tail forward velocity in [bl/s]. In contrast to control larvae (Rich media file 1), the rhythm is much slower, with alternating lower and higher maxima.

**Rich media file 3**: Animation of a sample *TH >ChR2-XXL* larva. The blue dot marks the position of the head. The filling indicates the absolute bending angle in [°]. At t = 30 s, light stimulation activates dopaminergic neurons; at t = 60 s, the light stimulation ends.

**Rich media file 4**: Animation of a sample *R53F07 >ChR2-XXL* larva. The blue dot marks the position of the head. The filling indicates the tail forward velocity in [bl/s]. At t = 30 s, light stimulation activates R53F07-neurons; at t = 60 s, the light stimulation ends.

### 9.2 Source files

**Figure 1-source file**: Source data underlying Figure 1-E1.

**Figure 2-source file**: Source data underlying Figures 2 and 2-E1, as well as precise sample sizes and results of statistical tests.

**Figure 3-source file**: Source data underlying Figure 3, as well as precise sample sizes and results of statistical tests.

**Figure 4-source file**: Source data underlying Figures 4 and 4-E1, as well as precise sample sizes and results of statistical tests.

**Figure 5-source file**: Source data underlying Figure 5, as well as precise sample sizes and results of statistical tests.

**Figure 6-source file**: Source data underlying Figure 6, as well as precise sample sizes and results of statistical tests.

**Figure 7-source file**: Source data underlying Figures 7 and 7-E1, as well as precise sample sizes and results of statistical tests.

**Figure 8-source file**: Source data underlying Figures 8 and 8-E1, as well as precise sample sizes and results of statistical tests.

**Figure 9-source file**: Source data underlying Figures 9 and 9-E1, as well as precise sample sizes and results of statistical tests.

**Figure 10-source file**: Source data underlying Figure 10, as well as precise sample sizes and results of statistical tests.

**Figure 11-source file**: Source data underlying Figure 11, as well as precise sample sizes and results of statistical tests.

**Figure 12-source file**: Source data underlying Figures 12 and 12-E1, as well as precise sample sizes and results of statistical tests.

